# An *in vivo* screen identifies NAT10 as a master regulator of brain metastasis

**DOI:** 10.1101/2024.08.20.608711

**Authors:** Jocelyn F Chen, Peng Xu, Wesley L Cai, Huacui Chen, Emily Wingrove, Wenxue Li, Giulia Biancon, Meiling Zhang, Amer Balabaki, Ethan D Krop, Yangyi Zhang, Mingzhu Yin, Toma Tebaldi, Jordan L Meier, Thomas F Westbrook, Stephanie Halene, Yansheng Liu, Don X Nguyen, Qin Yan

**Author notes:** These authors contributed equally: Jocelyn F Chen, Peng Xu. These authors contributed equally: Wesley L Cai, Huacui Chen, Emily Wingrove.

## Abstract

Metastasis is the major cause of cancer-related deaths. Emerging evidence has shown that epigenetic regulation plays a fundamental role in cancer metastasis. To better understand the epigenetic regulation of metastasis, we conducted an in vivo shRNA screen for vulnerabilities of brain metastasis and identified N-acetyltransferase 10 (NAT10) as a driver of brain metastasis. Knockdown of NAT10 significantly restrains cancer cell proliferation and migration *in vitro*, and tumor growth and brain metastasis *in vivo*. Structure-function analysis of NAT10 showed that its poorly characterized RNA helicase domain is critical for breast cancer cell growth *in vitro*, while its N-acetyltransferase domain is essential for primary tumor growth and brain metastasis *in vivo*. Integrative transcriptomic and proteomic analyses revealed key downstream effectors of NAT10, including PHGDH and PSAT1, two catalyzing enzymes for serine biosynthesis implicated in brain metastasis, and HSPA5, known to promote metastasis. We found that distant metastases of breast cancer, especially brain metastases express higher levels of NAT10, PHGDH, PSAT1, and HSPA5. Silencing PHGDH/PSAT1 or HSPA5 in metastatic breast cancer cells inhibits their ability to grow in the serine/glycine-limited condition or migrate, respectively, phenocopying the effects of NAT10 depletion. Moreover, NAT10 promotes the expression of PHGDH, PSAT1, and HSPA5 in its RNA helicase-dependent manner. These findings establish NAT10 as a master regulator of brain metastasis and shed light on the biological functions of its RNA helicase domain, nominating NAT10 as a target for treating metastatic diseases.

## Introduction

Metastasis is responsible for 90% of cancer-associated mortality^1^. Brain metastases, found in 15∼35% of patients with breast cancer and 16-27% of patients with lung cancer^2,3^, is associated with the worst outcome compared to metastasis to other organs^4^. Treatments for these patients, especially those with triple negative breast cancer (TNBC) brain metastasis, is limited. The median survival from the diagnosis of TNBC central nervous system metastases is only 4.9 months^5,6^. Thus, effective approaches to treat brain metastasis are urgently needed.

Epigenetic regulation has been found in all the steps of metastasis cascades^1^. Studies from our group and others have shown the profound effects of epigenetic alterations on the expression of cancer-essential genes or metastasis-promoting genes, which subsequentially modulates cancer cell proliferation, metastatic capability, and the adaptation to the distal metastatic sites^7–9^. However, the epigenetic dependencies of brain metastasis are largely unexplored.

Here, we constructed screens to specifically investigate the epigenetic dependencies of breast cancer brain metastasis (BCBM). We established NAT10 as a driver of BCBM with extensive evidence from *in vitro* experiments and preclinical studies. NAT10 is the only identified N-acetyltransferase that “writes” N4-acetylcytidine (ac4C) on RNA in mammals. It acetylates multiple species of RNA, including rRNA, tRNA, and possibly mRNA, and also appears to carry out lysine acetylation on proteins including histones^10–14^ and thus included in this screen. Emerging studies have revealed NAT10’s role in tumorigenesis and metastasis via its N-acetyltransferase function^15–18^. NAT10 also harbors an RNA helicase domain, the function of which is obscure in cancer biology. Here we show that mechanistically, NAT10 regulates PHGDH, PSAT1, and HSPA5 via its RNA-helicase domain, and these downstream effectors contribute collectively to proliferation and metastatic capability of breast cancer cells.

## Results

### An *in vivo* screen identifies NAT10 as a top drop-out hit of BCBM

To facilitate brain metastasis screen, we performed *in vivo* selection for brain metastatic derivatives of the MDA-MB-231-BrM2 (231-BrM2), a brain metastasis derivative of MDA-MB-231 TNBC cells^19^, and generated the MDA-MB-231-BrM3 (231-BrM3) derivative line with higher brain metastasis potential (Extended Data Fig. 1a). Compared with 231-BrM2, 231-BrM3 showed significantly increased brain metastasis ability as measured by bioluminescence *in vivo* and *ex vivo* (Extended Data Fig. 1b-c), making it an ideal tool to screen for regulators of brain metastasis.

To identify epigenetic dependencies for BCBM, we conducted parallel *in vivo* and *in vitro* functional screens using an inducible, barcoded shRNA library (Fig. 1a). This shRNA screening strategy was successfully employed in our previous breast cancer lung metastasis study^7^. Briefly, we compiled a list of epigenetic regulators based on the availability of small molecule inhibitors and the association between their expression and poor survival. We tested the knockdown efficiency of 326 shRNAs targeting 89 actionable epigenetic regulators and subcloned one shRNA with the best knockdown efficiency per target gene into the doxycycline (DOX) inducible and barcoded pINDUCER10 lentivirus backbone^7^. ShRNA against *BUD31*, which was previously shown to be essential for cell proliferation^20^, was included as the positive control, while shRNA against *CHEK1* as the negative control^7^. To enhance the representation of each factor in the screen, we generated six mini-pools by equally mixing 11-13 individual inducible epigenetic regulator knockdown cell lines with the positive and negative control cell lines mentioned above. Mini-pools were injected intracardially in mice or cultured on plates for *in vivo* or *in vitro* screens, respectively (Fig. 1a).

**Fig.1.**
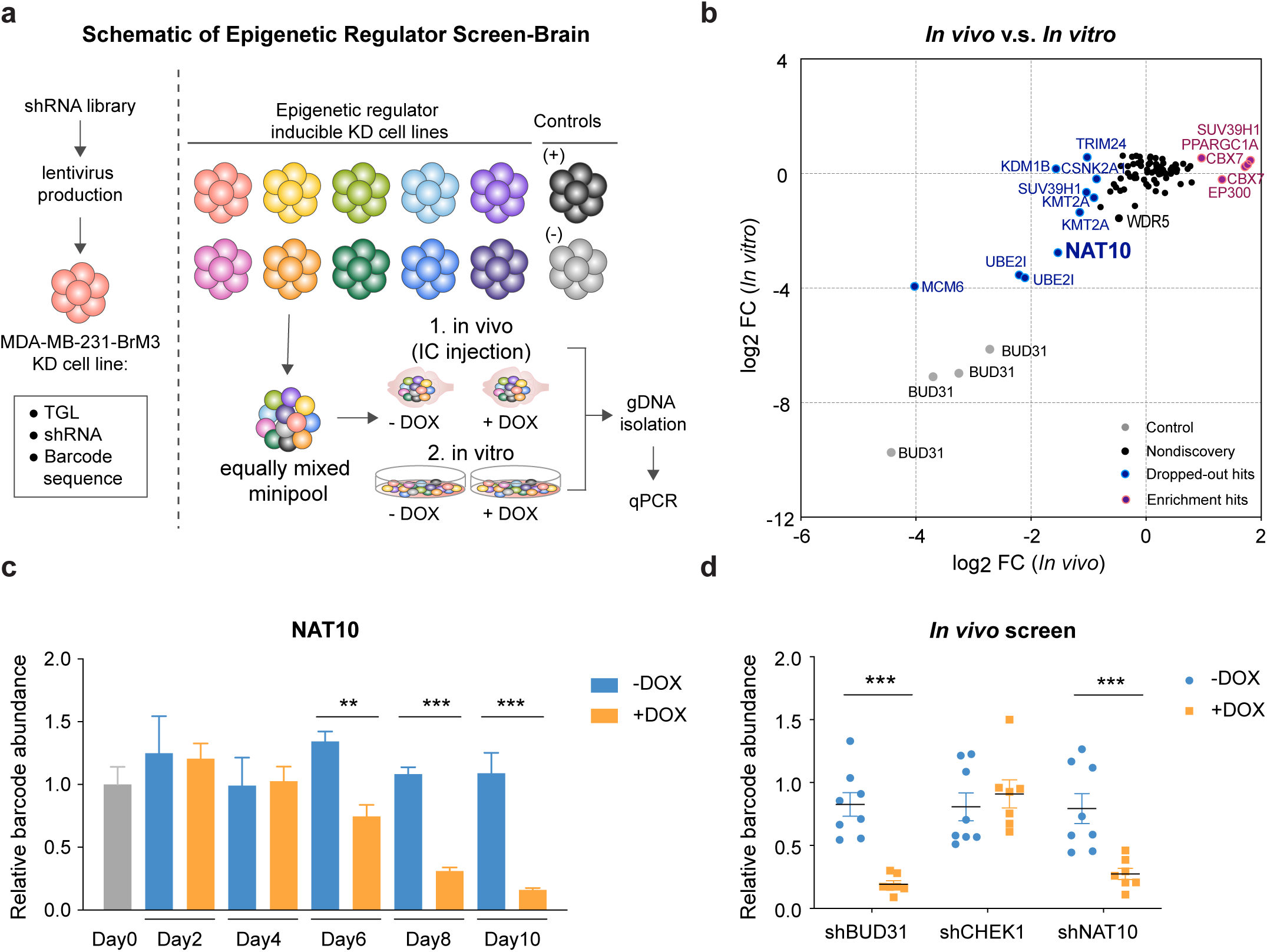
| *In vivo* and *in vitro* screens identified epigenetic dependencies of breast cancer brain metastasis. **a**, Schematic of shRNA screens for identifying the epigenetic dependencies of breast cancer brain metastasis. pGIPZ plasmid harboring barcode and hairpin targeting certain epigenetic factor was digested and sub-cloned into the pINDUCER10 plasmid, which contains HSV1-TK/GFP/Fluc (TGL) triple reporter gene. Epigenetic regulator inducible knockdown cell lines were equally mixed and injected into mice or cultured under control or doxycycline (1 µg/mL) treated condition. Brain (*in vivo*) and cells (*in vitro*) were harvested for gDNA and subjected to barcode qPCR as the screening output. **b**, Log_2_ (fold change) of the *in vivo* versus *in vitro* screen results for each epigenetic regulator. **c**, Relative abundance of barcode for shRNA against NAT10 in cells cultured with control or doxycycline treatment. **d**, Relative abundance of barcode for shRNA against NAT10, BUD31, and CHEK1 in brain tissue from control and doxycycline-treated mice. BUD31 and CHEK1 serve as positive and negative control, respectively. DOX, doxycycline. Significance in (**c**) and (**d**) were determined using unpaired student’s t-test. **, *P* < 0.01; ***, *P* < 0.001.

Out of 69 epigenetic regulators that were targeted in this screen, we identified 12 significant hits (*P* < 0.05, log_2_ (fold change) > 0.8 or < -0.8), including 8 drop-out and 4 enrichment hits (Fig. 1b and Extended Data Fig. 2a-h), with positive and negative controls showing corresponding phenotypes (Extended Data Fig. 2i-j). Some *in vivo* drop-out candidates also exhibited drop-out phenotype *in vitro*, including *MCM6*, *UBE2I*, and *NAT10* (Fig. 1b-d). MCM6 is an essential factor for DNA replication initiation^21^ and it was reported that MCM6 repression inhibits TNBC lung metastasis^22^. UBE2I/UBC9, the sole known E2 ubiquitin conjugating enzyme, was shown to promote growth of breast cancer in mice^23^. NAT10, the only ac4C N-acetyltransferase in mammals, acetylates multiple RNA species and proteins to modulate their stability and activity^12,13^. However, the roles of NAT10 in brain metastasis have not been investigated.

### Silencing of NAT10 inhibits breast cancer cell proliferation and migration

To dissect the functions of NAT10 in breast cancer, we generated two independent NAT10 inducible knockdown 231-BrM3 cell lines (Fig. 2a) and demonstrated that knockdown of NAT10 significantly decreased clonogenic ability of 231-BrM3 (Fig. 2b-c). Reduced cell proliferation can occur secondary to the disruption of cell cycle. To identify which phase within cell cycle progression is affected, we synchronized the cells at G2/M phase with Nocodazole and monitored the percentage of cells entering each phase of cell cycle after release from G2/M arrest (Extended Data Fig. 3a). NAT10 knockdown cells exhibited slower progression from G2/M to S phase (Fig. 2d and Extended Data Fig.3b). Consistently, NAT10 knockdown cells showed a decreased BrdU incorporation rate compared to control cells (Fig. 2e), indicating that NAT10 is critical for cells to enter S phase. Importantly, knockdown of NAT10 downregulated Cyclin D1 (Fig. 2a), a key regulator of the cell cycle progression. Reduced clonogenic ability could also be due to enhanced cell death, we therefore assessed whether NAT10 knockdown results in apoptosis in 231-BrM3 cells. Only less than one percent of cells in NAT10 knockdown and control groups undergo apoptosis (Extended Data Fig. 3c-d), suggesting that knockdown of NAT10 does not induce apoptosis.

**Fig.2.**
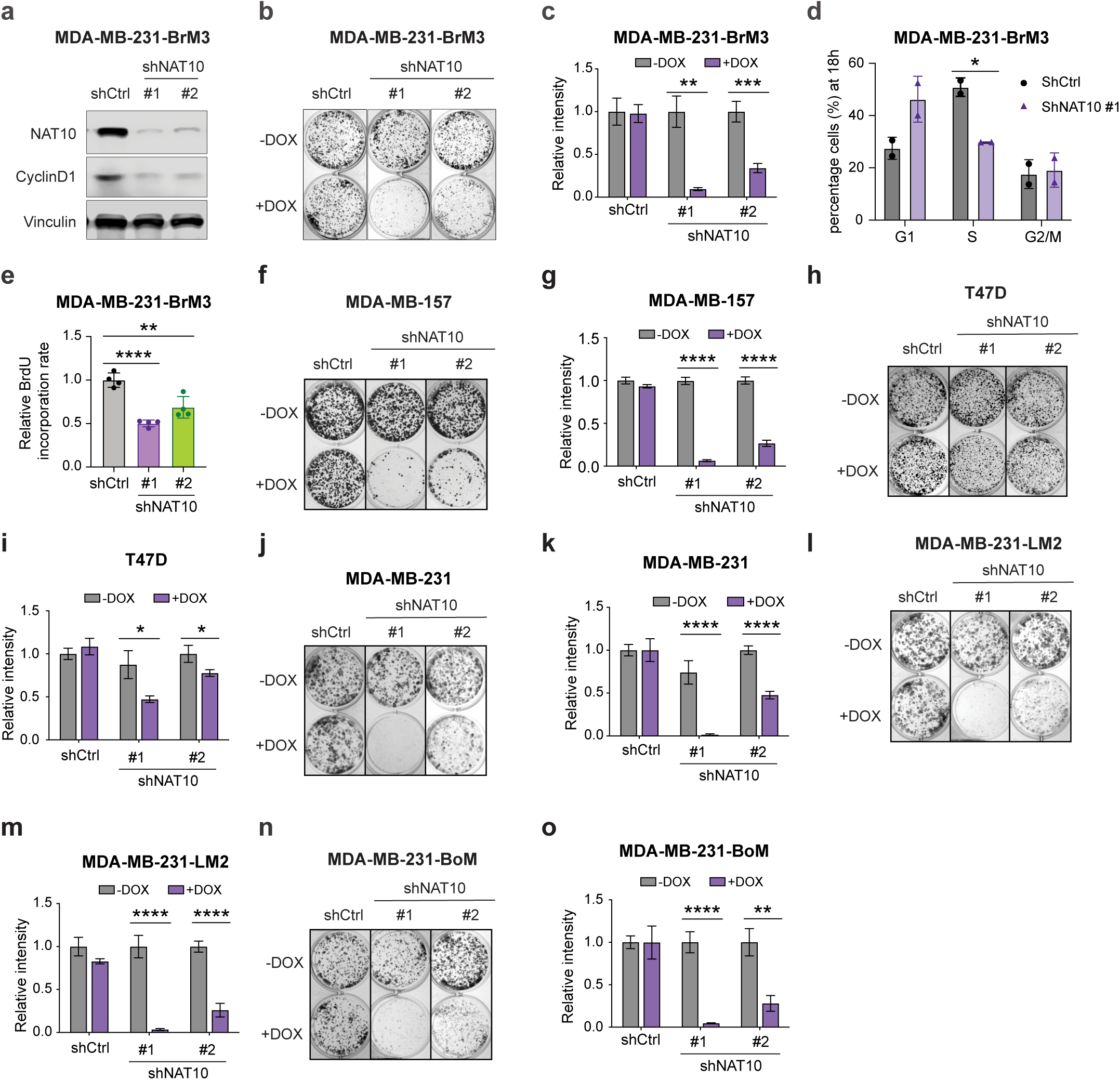
| NAT10 promotes breast cancer proliferation across subtypes and metastatic organotropisms. **a**, Western blot of indicated proteins in 231-BrM3 cells harboring inducible control or NAT10 targeting shRNAs (shNAT10 #1 and shNAT10 #2) after 3 days of doxycycline (1 µg/mL) induction. **b-c**, Colony formation assays of 231-BrM3 cells with NAT10 knockdown or control after 9 days of either control or doxycycline (1 µg/mL) treatment. Representative images (**b**) and quantification (**c**) are shown. **d**, Percent of cells in G1, S, and G2/M phase from indicated cells fixed at 18 hours post-refeeding. **e**, BrdU incorporation assay of 231-BrM3 cells after 3 days of doxycycline (1 µg/mL) induction. **f-o**, Colony formation assays of indicated cells after 9 days of either control or doxycycline (1 µg/mL) treatment. Representative images (**f**, **h**, **j**, **l**, **n**) and quantification (**g**, **i**, **k**, **m**, **o**) are shown. Significance in (**c-e**, **g**, **i**, **k**, **m**, **o**) was determined using unpaired student’s t-test. *, *P* < 0.05; **, *P* < 0.01; ***, *P* < 0.001; ****, *P* < 0.0001.

### NAT10 promotes proliferation across breast cancer subtypes and metastatic organotropisms

We further asked whether NAT10-regulated cell proliferation is a general phenomenon in other breast cancer subtypes and MDA-MB-231 derivatives with distinct organotropism. We examined effects of NAT10 knockdown on clonogenic ability in an additional TNBC cell line MDA-MB-157 and an estrogen receptor (ER) positive breast cancer cell line T47D. Knockdown of NAT10 impaired the clonogenic ability of MDA-MB-157 and T47D (Fig. 2f-i and Extended Data Fig. 3e-f). It is noteworthy that the effect of NAT10 loss on MDA-MB-157 and 231-BrM3 is stronger than that on T47D, suggesting that TNBC cells may be more dependent on NAT10 than ER positive breast cancer cells.

To address whether NAT10 is essential for the growth for TNBC cells with distinct organotropism, we turned to two other organotropic derivatives of MDA-MB-231, MDA-MB-231-LM2 (231-LM2) and MDA-MB-231-BoM (231-BoM). Having been selected *in vivo* similarly with 231-BrM3, 231-LM2 and 231-BoM cells preferentially metastasize to lung and bone, respectively^19,24^. We depleted NAT10 with two independent shRNAs in parental MDA-MB-231, 231-LM2, 231-BoM (Extended Data Fig. 3g-h) and found that knockdown of NAT10 significantly impaired colony formation in parental MDA-MB-231, 231-LM2, and 231-BoM to similar degrees as in 231-BrM3 cells (Fig. 2j-o and Extended Data Fig. 3i-k). These results demonstrate NAT10 regulates cell proliferation independent of their metastatic organotropism.

### NAT10 is required for breast tumor growth and metastasis

To test whether knockdown of NAT10 affects orthotopic tumor growth *in vivo*, we injected 231-BrM3 cells with or without NAT10 knockdown to the 4^th^ mammary fat pads in mice. A significant decrease of bioluminescence signal in mammary glands was observed in NAT10 knockdown group compared to control group (Fig. 3a). At the end point (day 58), the tumor size and weight in NAT10 knockdown group were significantly reduced compared to the control tumors (Fig. 3b-d).

**Fig.3.**
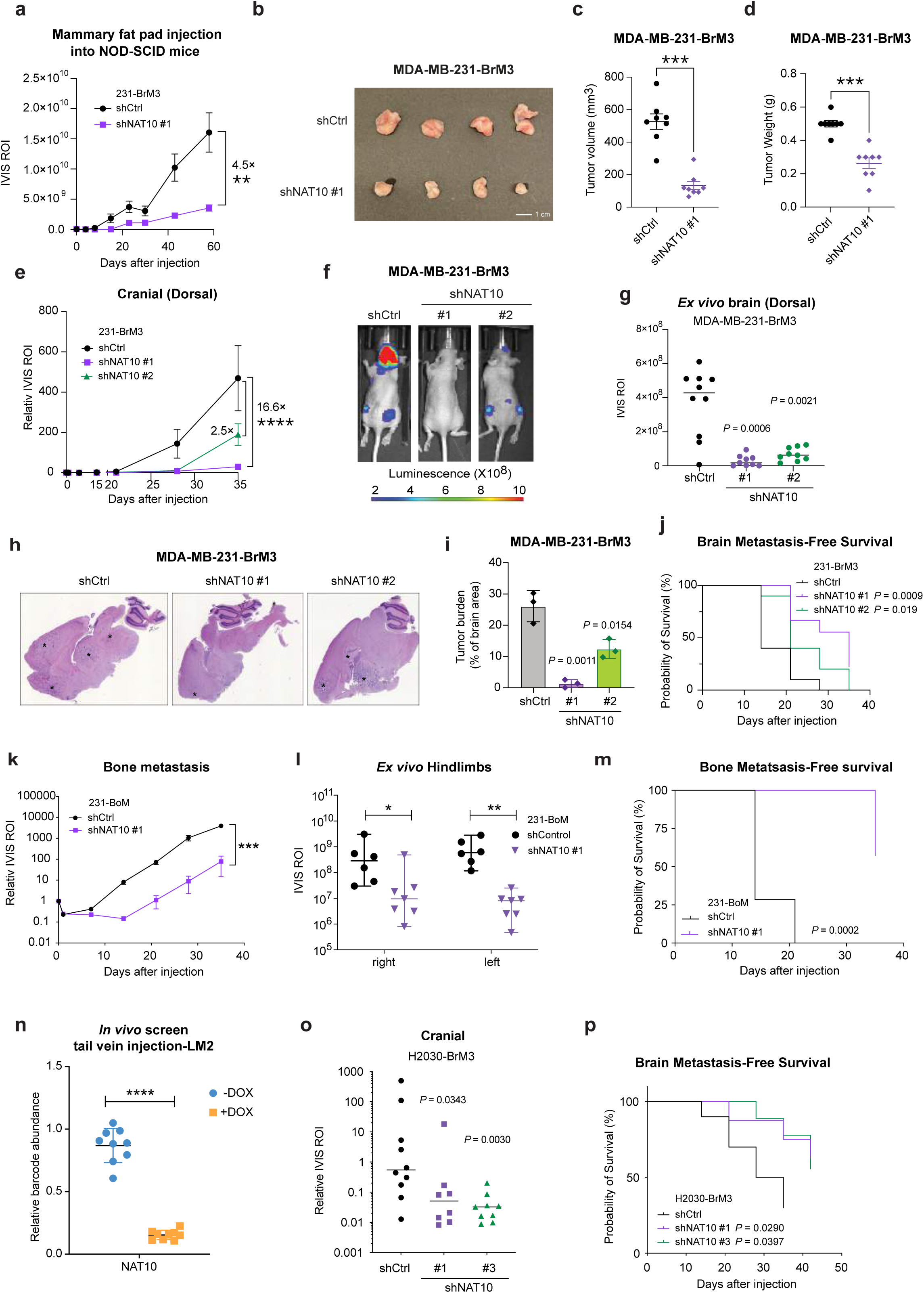
| NAT10 is essential for primary breast tumor growth and brain metastasis. **a**, Bioluminescence signals of mice injected into the 4^th^ mammary fat pad with 231-BrM3 cells harboring inducible control or shNAT10 #1. The data represent average ± SEM (Unpaired student’s t-test; **, *P* < 0.01). **b**, Representative images of primary tumors from mice in (**a**) at day 58. **c-d**, Quantification of primary tumor volume (**c**) and weight (**d**) from mice in (**a**) at day 58. Each dot represents one mouse (Unpaired two-tailed Mann-Whitney test; ***, *P* < 0.001). **e**, Normalized bioluminescence signals of brain metastases of mice injected intracardially with 231-BrM3 cells harboring inducible control, shNAT10 #1, or shNAT10 #2 and kept under doxycycline chow. The data represent average ± SEM (Unpaired student’s t-test; ****, *P* < 0.0001). **f**, Representative bioluminescence images of mice in (**e**) at day 28. **g**, Quantification of *ex vivo* bioluminescence signals of brain tissues from mice in (**e**) at day 35 post-injection (Unpaired two-tailed Mann-Whitney test). Each dot represents one mouse. **h**, Representative hematoxylin and eosin (H&E) stain of sagittal brain tissue slides from shCtrl and shNAT10 groups. Brain metastases were marked with *. **i**, Quantification of tumor burden with ImageJ. n = 3 for each group (Two-tailed student’s t-test). **j**, Kaplan-Meier plot of brain metastasis-free survival in (**e**). shCtrl (n = 10), shNAT10 #1 (n = 9), and shNAT10 #2 (n = 10) were analyzed (Log rank Mantel-Cox test). **k**, Normalized bioluminescence signals of bone metastasis of mice injected intracardially with 231-BoM cells harboring inducible control or shNAT10 #1 and kept under doxycycline chow. The data represent average ± SEM (Unpaired student’s t-test; ***, *P* < 0.001). **l**, Quantification of *ex vivo* bioluminescence signals of hindlimbs (left and right) from mice in (**k**) at day 35 post-injection. Each dot represents one animal (Unpaired two-tailed Mann-Whitney test). **m**, Kaplan-Meier plot of bone metastasis-free survival in (**k**) (Log rank Mantel-Cox test). **n**, Relative abundance of barcode for shRNA against NAT10 in lung tissue from control and doxycycline-treated mice injected intravenously with 231-LM2 cells. Each dot represents one mouse (Unpaired student’s t-test; ****, *P* < 0.0001). **o,** Quantification of brain metastasis burden from mice injected intracardially with H2030-BrM3 cells at day 35 (Unpaired two-tailed Mann-Whitney test). **p**, Kaplan-Meier plot of brain metastasis-free survival in (**o**) (Log rank Mantel-Cox test).

To address whether NAT10 is required for BCBM, we injected 231-BrM3 cells with or without NAT10 knockdown through the left ventricle of mice. We monitored brain metastatic colonization and outgrowth over 35 days and observed a significant impairment on brain colonization in NAT10 knockdown group (Fig. 3e-g). At the end point (day 35), the average brain metastatic burden in mice with stronger NAT10 knockdown efficacy (shRNA #1, Fig. 2a) was 16.6-fold lower than that in mice injected with 231-BrM3 control cells (Fig. 3e-f). Notably, the difference in brain metastasis exceeds the difference in mammary tumor growth (comparing Fig. 3e with Fig. 3a), implying NAT10 has brain metastasis specific roles. Consistently, mice from NAT10 knockdown group showed decreased brain metastases burden as quantified via *ex vivo* brain luminescent signal at the endpoint (Fig. 3g) and as evident by the hematoxylin and eosin (H&E) staining of brain tissue sections (Fig. 3h-i).

As a small proportion of 231-BrM3 cells can metastasize elsewhere in addition to the brain, we monitored whole-body metastasis and observed decreased overall metastatic burden in NAT10 knockdown groups (Extended Data Fig. 4a). Mice injected with NAT10-depleted 231-BrM3 cells showed delayed development of brain metastases and extracranial metastases compared to the control mice (Fig. 3j and Extended Data Fig. 4b). These results suggest that NAT10 is a critical factor for the development of breast cancer metastasis to the brain and other organs.

### NAT10 is an essential factor in multiple metastasis settings

Since NAT10 depletion also impaired whole-body metastasis burden (Extended Data Fig. 4a) and knockdown of NAT10 in both 231-BoM and 231-LM2 led to significant decrease in cell growth (Fig. 2k-p and Extended Data Fig. 3j-l), we asked whether NAT10 affects breast cancer metastasis to other organs such as bone and lungs. We injected the 231-BoM cells with or without NAT10 knockdown intracardially and monitored bone metastasis colonization and outgrowth especially at the hindlimb area. We observed a significant impairment on bone colonization in NAT10 knockdown group (Fig. 3k). Consistently, mice from NAT10 knockdown group also showed decreased hindlimb metastases burden in both sides from the quantification of *ex vivo* bone luminescent signal at the end point (Fig. 3l). Mice injected with NAT10-depleted 231-BoM cells showed delayed development of bone metastases and non-bone metastases compared to the control mice (Fig. 3m and Extended Data Fig. 4c). In addition to bone metastasis, we revisited our previous screening results of breast cancer lung metastasis^7^ and found that 231-LM2 cells with NAT10 ablation was significantly dropped-out in the lungs (Fig. 3n).

Among the brain metastasis cases, metastases originating from lung cancer account for the highest percentage^25^. To test whether NAT10 is required for brain metastasis by lung cancer, we utilized H2030-BrM3 cells, brain metastatic derivatives of the H2030 non-small cell lung cancer cell line harboring the *KRAS^G12C^*mutation. We depleted NAT10 using shRNA (Extended Data Fig. 4d) and injected H2030-BrM3 with or without NAT10 knockdown intracardiacally into the mice. Knockdown of NAT10 significantly decreased the brain metastasis ability of H2030-BrM3 (Fig. 3o). Additionally, mice injected with NAT10-depleted H2030-BrM3 cells showed delayed development of brain metastases and extracranial metastases compared to the control mice (Fig. 3p and Extended Data Fig. 4e). Taken together, we conclude that NAT10 is a driver of breast and lung cancer metastasis to multiple organs.

### The RNA helicase function of NAT10 is critical for breast cancer cell growth

We next sought to examine which molecular function of NAT10 is required for breast cancer cell growth. To this end, we performed structural functional analysis of NAT10 by constitutively expressing shRNA-resistant NAT10 wild type as well as three NAT10 mutants (G641E, acetyltransferase dead; K290A, helicase dead; K426R, mutant of NAT10 autoacetylation site) ^26–28^ with N-terminal Flag-tag in the NAT10 knockdown cell line (Fig. 4a). We optimized the condition for DOX induction so that the expression levels of ectopic NAT10 (wild type or mutants) were comparable to endogenous NAT10 in the indicated cell lines, whereas endogenous NAT10 was significantly depleted (Fig. 4b).

**Fig.4.**
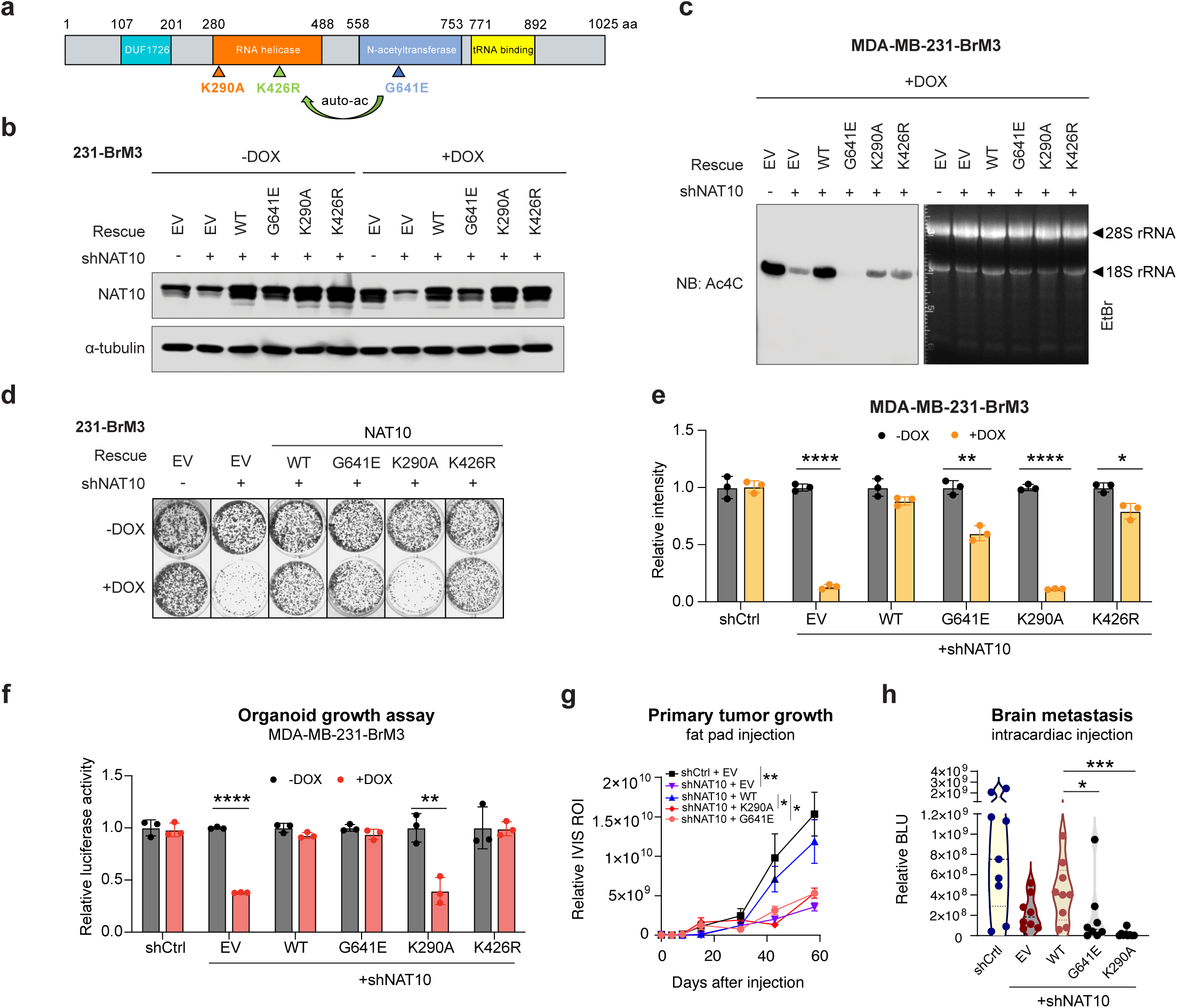
| RNA helicase and N-acetyltransferase domains are required for NAT10-mediated phenotypes. **a**, Schematic of NAT10 with indicated domains and mutation sites. G641E, acetyltransferase dead mutant; K290A, helicase dead mutant; K426R, NAT10 autoacetylation site mutant. **b**, Western blot of indicated proteins in 231-BrM3 cells with inducible control or shNAT10, followed by rescuing with empty vector (EV), shRNA-resistant wild type (WT) NAT10 or NAT10 mutants. Cells were collected after 3 days of control or doxycycline (1 µg/mL) induction. **c**, Immuno-Northern blots of ac4C modification in total RNA extracted from indicated genetically manipulated 231-BrM3 cells. Cells were induced with doxycycline (1 µg/mL) for 5 days. **d-e**. Colony formation assays of indicated genetically manipulated 231-BrM3 cells after 9 days of either control or doxycycline (1 µg/mL) treatment. Representative images (**d**) and quantification (**e**) are shown. **f**, Organoid growth assays of indicated genetically manipulated 231-BrM3 cells after 12 days of either control or doxycycline (1 µg/mL) treatment. The 3D growth of cells was determined by *in vitro* luciferase activity and normalized to control. **g**, Bioluminescence signals of mice injected into the 4^th^ mammary fat pad with indicated genetically manipulated BrM3 cells. The data represent average ± SEM. **h**, Quantification of bioluminescence signals of brain metastases in mice injected intracardially with indicated genetically manipulated 231-BrM3 cells at day 35. Each dot represents one animal. Unpaired student’s t-test was used in (**e**-**g**), while unpaired two-tailed Mann-Whitney test was used in (**h**). *, *P* < 0.05; **, *P* < 0.01; ***, *P* < 0.001; ****, *P* < 0.0001.

As NAT10 is capable of acetylating several reported RNAs and proteins, we assessed whether such acetyltransferase activity is active in 231-BrM3 and whether such modification contributes to the observed growth phenotype. We identified a strong band at the position of 18S rRNA, a known RNA substrate of NAT10^11^, with ac4C antibody in immune-northern blot, while the signal decreased when NAT10 was inducibly depleted (Extended Data Fig. 5a). In the time course experiment, the ac4C on 18S rRNA decreased over time (Extended Data Fig. 5b). We also observed light NAT10-dependent smears occurring in the size range expected for poly(A) RNAs (Extended Data Fig. 5a-b). Similarly, NAT10-dependent 18S rRNA acetylation was detected in H2030-BrM3 cells (Extended Data Fig. 5c-d). We then examined the RNA acetylation in NAT10 rescued cell lines and observed that re-expressing wild type NAT10 restored the ac4C modification on 18S rRNA (Fig. 4c). The 18S rRNA acetylation was completely abolished when the G641E mutant was introduced, indicating the G641E mutant serves a dominant negative mutant, consistent with the notion that this residue is critical for NAT10’s acetyltransferase activity^26^. Interestingly, the K290A and K426R mutants were unable to fully rescue 18S rRNA acetylation (Fig. 4c), indicating these mutants also have impaired acetyltransferase activity. Together, these results confirmed that NAT10 is responsible for RNA ac4C in breast and lung cancer cells.

To determine which function of NAT10 contributes to cell growth, we performed colony formation assays using the rescued cell lines treated with DOX or DMSO. We observed that re-expressing wild type NAT10 rescued the growth phenotype by around 85%, a similar result was observed when we re-expressed the K426R mutant (Fig. 4d-e). Surprisingly, G641E mutant could partially rescue the growth to more than 50%. In contrast, K290A mutant could not rescue the growth at all (Fig. 4d-e). To examine the roles of NAT10 in a more physiologically relevant setting, we performed 3D organoid growth assay by culturing 231-BrM3 cells in suspension with 5% Matrigel. Again, we observed that NAT10 K290A mutant re-expression did not rescue the 3D growth of 231-BrM3 cells while wild type, G641E, and K426R mutants did (Fig. 4f). These results suggest that the RNA helicase function, but not the acetyltransferase activity or autoacetylation of NAT10, is most critical for breast cancer cell growth.

### The N-acetyltransferase function of NAT10 is essential for breast cancer growth and brain metastasis *in vivo*

To further explore which function of NAT10 is essential for the growth of 231-BrM3 *in vivo*, we injected the same panel of genetically manipulated 231-BrM3 cells into the 4^th^ mammary fat pad of NOD-SCID mice to evaluate their effects on primary tumor growth. As expected, wild type NAT10 was able to rescue the primary tumor growth (Fig. 4g). However, neither the G641E mutant nor the K290A mutant was able to rescue tumor growth (Fig. 4g). Considering the K290A mutant is also defective in 18S rRNA acetylation (Fig. 4c), these results suggest that the acetyltransferase function is critical for breast tumor growth *in vivo*.

When injected intracardiacally into the mice, NAT10 K290A and G641E mutant-rescued cells also exhibited impaired brain metastasis colonization and outgrowth as compared to the wild type NAT10 rescued cells (Fig. 4h). The K290A mutant is more defective than G641E and vector control (Fig. 4h), suggesting that it has dominant negative effect. These results showed that the N-acetyltransferase function is critical for brain metastasis *in vivo,* while the RNA helicase function also contributes to brain metastasis *in vivo*. Collectively, our results demonstrated that the acetyltransferase function is required for breast tumor growth and brain metastasis *in vivo*.

### NAT10 promotes PHGDH, PSAT1, and HSPA5 expression in an RNA helicase domain-dependent manner

To elucidate the molecular effects and biological pathways of NAT10 depletion, we performed RNA sequencing (RNA-Seq) analysis of NAT10-depleted and control 231-BrM3 cells. The most enriched Biological Process (BP) terms in Gene Ontology (GO) analysis of differentially expressed genes included regulation of cell proliferation, cell migration, L-serine biosynthetic process, and cellular response to glucose stimulus (Extended Data Fig. 6a). To narrow down the downstream effector candidates, we also performed data-independent acquisition mass spectrometry (DIA-MS) analysis^29^ of NAT10 knockdown and control 231-BrM3 cells. The most enriched BP terms among the differentially expressed proteins included terms related DNA replication, translation, and L-serine biosynthetic process (Extended Data Fig. 6b). When overlapping differentially expressed candidates from RNA-Seq and DIA-MS, we identified 11 potential downstream factors which showed consistent changes (Fig. 5a).

**Fig.5.**
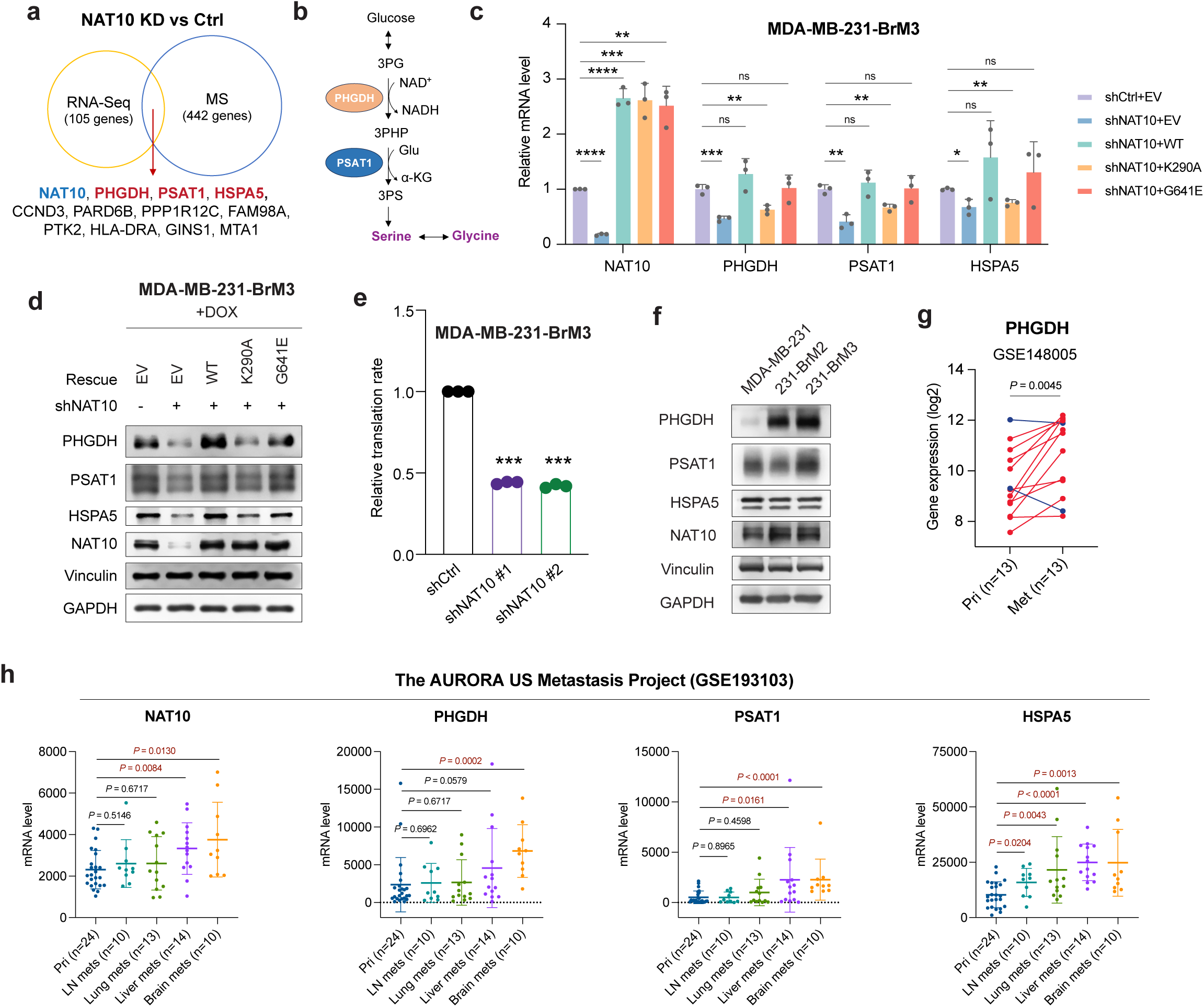
| NAT10 promotes PHGDH, PSAT1, and HSPA5 expression in an RNA helicase dependent manner. **a**, The overlap of differentially expressed genes in RNA-Seq and differentially expressed proteins in data independent acquisition mass spectrometry (DIA-MS) of 231-BrM3 cells with NAT10 knockdown (shNAT10 #1) versus control cells (shCtrl). For both RNA-Seq and DIA-MS, the differentially expressed candidates were defined as *P* < 0.5 and log_2_ (Fold Change) > 0.3 or < -0.3. **b**, Glucose derived L-Serine biosynthesis pathway. 3PG, 3-phosphoglycerate; 3-PHP, 3-phosphohydroxypyruvate; 3PS, 3-phosphoserine; Glu, glutamate. **c-d**, Relative mRNA levels (**c**) and protein levels (**d**) in 231-BrM3 cells with indicated genetic manipulations. **e**, The relative translation rate of 231-BrM3 with or without NAT10 knockdown. **f**, Western blots of indicated proteins in parental MDA-MB-231, 231-BrM2, and 231-BrM3 cell lines. **g**, PHGDH mRNA levels in 13 paired primary breast tumors and metastatic breast tumors from our previous dataset (GSE148005). In total, 11 out of 13 metastatic breast tumors show higher PHGDH expression than it in primary tumors (lines in red). **h**, NAT10, PHGDH, PSAT1, and HSPA5 mRNA levels in primary breast tumors and their matched metastases in The AURORA US Metastasis Project (GSE193103). Pri, primary breast tumors; LN mets, lymph node metastasis. *P* values are marked in red if lower than 0.05. Unpaired student’s t-test was used in (**c** and **e**), while paired student’s t-test was used in (**g** and **h**). ns, not significant; *, *P* < 0.05; **, *P* < 0.01; ***, *P* < 0.001; ****, *P* < 0.0001.

Among the 11 potential downstream effectors of NAT10, 3-phosphoglycerate dehydrogenase (PHGDH) and phosphoserine aminotransferase 1 (PSAT1) are catalyzing enzymes in the glucose-derived serine/glycine synthesis pathway (Fig. 5b), in which PHGDH oxidizes the glycolytic intermediate 3-phosphoglycerate (3PG) to 3-phosphohydroxypyruvate (3-PHP), which is subsequently converted to 3-phosphoserine (3PS) by PSAT1 and eventually to serine by phosphoserine phosphatase (PSPH). Serine is catabolized to generate glycine by mitochondrial serine hydroxymethyltransferase 2 (SHMT2) or cytosolic SHMT1^30,31^. Serine and glycine are essential resources for protein synthesis and crucial for cell proliferation^30,31^. Importantly, PHGDH-mediated L-serine biosynthesis is crucial for cancer cells to survive the serine and glycine-limited microenvironment^32,33^. In addition, heat-shock protein 5 (HSPA5) has been identified as a poor prognosis marker for in breast cancer and shown to promote breast cancer metastasis^34^.

To validate whether NAT10 regulates PHGDH, PSAT1, and HSPA5 in 231-BrM3, we turned to NAT10 knockdown cell lines and confirmed that their mRNA levels significantly decreased when NAT10 was depleted (Fig. 5c). We further interrogated which function of NAT10 is critical for this regulation by using NAT10 wild-type, K290A, or G641E mutant-rescued cell lines. PHGDH, PSAT1, and HSPA5 expression could be rescued by wild-type NAT10 (Fig. 5c). Intriguingly, their expression could be rescued by G641E mutant as well, but not K290A mutant (Fig. 5c). This RNA-helicase dependent regulation of NAT10 could also be validated at the protein level (Fig. 5d). These data suggest that NAT10 promotes PHGDH, PSAT1, and HSPA5 expression in an RNA-helicase dependent manner. We further measured protein translation rate when NAT10 was depleted and showed that silencing NAT10 impaired global translation rate (Fig. 5e), consistent with the translational regulation role of NAT10.

### NAT10, PHGDH, PSAT1, and HSPA5 are highly expressed in brain metastases

We examined the expression levels of NAT10 and its downstream factors in parental MDA-MB-231 cell line and its brain metastasis derivatives. 231-BrM3 expressed the highest levels of NAT10, PHGDH, and PSAT1, but not HSPA5 (Fig. 5f and Extended Data Fig. 6c). Both mRNA and protein levels of PHGDH increased proportionally to the increasing brain metastatic ability of 231-BrM2 and 231-BrM3 (Fig. 5f and Extended Data Fig. 6c). Consistently, RNA sequencing results showed upregulation of PHGDH in brain metastasis derivatives, including 231-BrM2 in our previous RNA-seq dataset (GSE138122) (Extended Data Fig. 6d)^35^ and MDA-MB-231 brain metastasis variant (231-BR) in a publicly available RNA-seq dataset (GSE183862) (Extended Data Fig. 6e)^36^.

To examine the relevance of our findings in human breast tumors, we analyzed our previously published dataset containing 13 primary breast cancer tissues and their paired metastases (GSE148005)^8^. PHGDH was highly expressed in 11 out of 13 metastases compared to the paired primary breast cancer tissues (Fig. 5g). We further investigated their expression in The AURORA US Metastasis Project (GSE193103)^37^, a larger cohort with paired breast cancer metastases and primary tumors. NAT10, PHGDH, PSAT1, and HSPA5 were highly expressed in metastases compared with the primary breast tumors (Extended Data Fig. 6f). Importantly, among all main metastasis sites of breast cancer, brain metastases exhibited the highest levels of NAT10, PHGDH, PSAT1, and HSPA5, followed by liver metastases (Fig. 5h).

### Depletion of PHGDH/PSAT1 sensitizes cancer cells to serine/glycine deprivation

PHGDH and PSAT1 are essential enzymes for glucose-derived serine/glycine biosynthesis pathway; by utilizing glucose, metastatic breast cancer cells synthesize serine/glycine to survive the amino acid–limited brain microenvironments^31–33^. To investigate the proliferation rate of metastatic breast cancer cells in brain microenvironment, we made the cerebrospinal fluid (CSF)-Like medium that contains limited serine/glycine^33^ and a harsher medium, i.e. w/o-Ser/Gly that does not contain serine/glycine at all, compared to the DMEM-Like medium that contains high levels of serine/glycine (Fig. 6a).

**Fig. 6.**
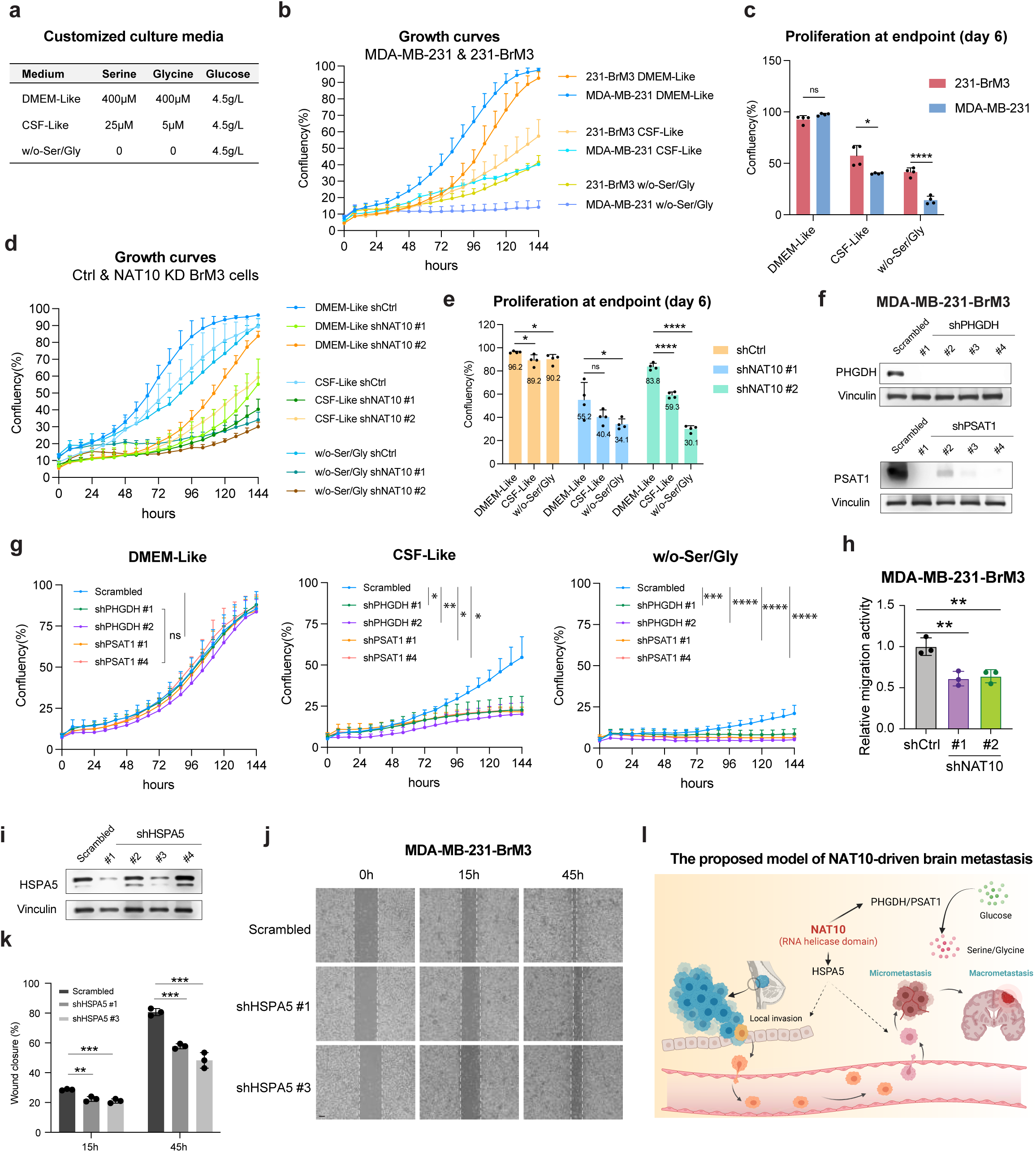
| NAT10-regulated PHGDH, PSAT1, and HSPA5 contribute to the characteristics of metastatic breast cancer cells. **a**, The concentration of serine, glycine, and glucose in DMEM-Like, CSF-Like, and w/o-Ser/Gly media. **b-c**, Growth curves of 231-BrM3 and MDA-MB-231 cells in customized culture media (**b**) and the confluency at the endpoint (day 6) (**c**). **d-e**, Growth curves of 231-BrM3 cells with or without NAT10 knockdown in customized culture media (**d**) and the confluency at the endpoint (day 6) (**e**), with average confluency labeled. **f**, Western blots analysis of 231-BrM3 cells with scrambled control and shRNA against *PHGDH* or *PSAT1*. **g**, Growth curves of control cells or 231-BrM3 cells with PHGDH or PSAT1 knockdown in customized culture media. **h**, *In vitro* cell migration assay of 231-BrM3 cells after 3 days of doxycycline (1 µg/mL) induction. **i**, Western blots analysis of 231-BrM3 cells with scrambled control and shRNA against *HSPA5*. **j-k**, Representative images (**j**) and quantification (**k**) of wound healing assay of 231-BrM3 cells with or without HSPA5 knockdown. **l**, A model depicting the roles of NAT10 in brain metastasis, created with BioRender. Unpaired student’s t-test was used in (**c**, **e, h,** and **k**). ns, not significant; *, *P* < 0.05; **, *P* < 0.01; ***, *P* < 0.001; ****, *P* < 0.0001.

When cultured in DMEM-Like medium, 231-BrM3 and parental MDA-MB-231 proliferated similarly, with no significant difference at the endpoint (day 6) (Fig. 6b-c). However, 231-BrM3 cells proliferated faster than MDA-MB-231 cells in CSF-Like and w/o-Ser/Gly media (Fig. 6b-c). These results indicated that 231-BrM3 cells, which show much higher expression of PHGDH compared with parental MDA-MB-231 cells (Fig. 5f), survive better in serine/glycine limited or depleted environment.

To interrogate whether NAT10 contributes to the better survival of 231-BrM3 cells in serine/glycine limited or depleted media, we cultured 231-BrM3 cells with or without NAT10 depletion in the three customized media and monitored their growth curves. NAT10 depletion led to bigger difference in proliferation in CSF-Like or w/o-Ser/Gly medium than in DMEM-Like medium (Fig. 6d-e).

To further explore the roles of PHGDH and PSAT1 in the proliferation of 231-BrM3 in brain microenvironment, we depleted them using multiple shRNAs (Fig. 6f) and monitored their proliferation rate in above-mentioned three culture media. As expected, significant difference in proliferation of these cells was only observed CSF-Like medium and w/o-Ser/Gly media, but not in DMEM-like medium (Fig. 6g). Collectively, these results showed that NAT10 and its downstream factors PHGDH/PSAT1 are crucial for the proliferation of metastatic breast cancer cells in the serine/glycine-limited environment.

### Depletion of HSPA5 attenuates the migration ability of metastatic breast cancer cells

Previous study reported that increased HSPA5 could enhance the migration/invasive ability of breast cancer cells^34^. Considering NAT10 is the upstream regulator of HSPA5, we asked whether NAT10 has a role in migration ability of 231-BrM3 cells and observed decreased migration activity upon NAT10 depletion (Fig. 6h).

We further depleted HSPA5 in 231-BrM3 (Fig. 6i) and revealed that knockdown of HSPA5 attenuated the migration ability of 231-BrM3 cells (Fig. 6j-k), phenocopying the effect of NAT10 depletion and consistent with previous report^34^. HA15, a potent and specific HSPA5 inhibitor that inhibits the ATPase activity of HSPA5^38^, also decreased the migration ability of 231-BrM3 in a dose dependent manner (Extended Data Fig. 7a-b), without changing HSPA5 protein level (Extended Data Fig. 7c). In contrast, HSPA5 depletion has no significant effect on colony formation ability (Extended Data Fig. 7d-e). Taken together, our results demonstrate that PHGDH, PSAT1, and HSPA5 contribute collectively to NAT10-mediated proliferation and metastatic capacity (Fig. 6l).

## Discussion

In the current study, we performed *in vivo* and *in vitro* screens of epigenetic dependencies for BCBM and identified NAT10 as a top drop-out hit. We established NAT10 as a driver of BCBM through extensive *in vitro* and *in vivo* interrogations. *In vitro* assays showed that the RNA helicase domain, but not N-acetyltransferase domain, is the most crucial domain for the growth of metastatic breast cancer cells. Interestingly, *in vivo* data revealed that N-acetyltransferase domain is essential for tumor growth and brain metastasis. Furthermore, we demonstrated that NAT10 regulates the expression levels of PHGDH, PSAT1, and HSPA5 in an RNA-helicase dependent manner. NAT10 promotes PHGDH and PSAT1 expression to ensure a better survival in serine/glycine limited environment and HSPA5 expression to increase the migration ability, suggesting NAT10 functions as a master driver of brain metastasis that participates in multiple steps of metastatic cascade (Fig. 6l).

Our rescue experiments confirmed that G641 is the critical residue for acetyltransferase activity. In addition, we found the mutation of K290A impaired its ability to acetylate 18S rRNA (Fig. 4c), suggesting the ATPase activity of the RNA helicase is important for NAT10 to perform RNA acetylation. K426 is reported as an autoacetylation site of NAT10 and K426 acetylation is required for NAT10 to activate rRNA transcription^28^. Here, we uncovered that K426 is crucial for the acetyltransferase activity of NAT10 (Fig. 4c), adding new values to this overlooked site. The mechanism by which NAT10 autoacetylation contributes to its acetyltransferase activity requires further study. Our functional studies suggest the RNA acetyltransferase function of NAT10 is less important for breast cancer cell proliferation *in vitro* (Fig. 4d-f). Our results are consistent with the previous findings that yeast Kre33 and human NAT10 acetylate 18S rRNA and tRNA; however, rRNA acetylation is dispensable for yeast cell growth^11^. In contrast, we also found that the N-acetyltransferase function of NAT10 is required for breast tumor growth and metastasis in mice. These results indicate that NAT10 N-acetyltransferase inhibitors could be used to treat breast cancer metastasis.

We also show that breast cancer cell growth and NAT10 downstream genes PHGDH, PSAT1 and HSPA5 were regulated by NAT10 in its RNA helicase activity dependent manner. RNA helicases are a large class of enzymes that play pleotropic functions at multiple steps of the gene expression processes and the two largest families in humans are DEAD box (DDX) or DEAH box (DHX) proteins^39^. Structurally, the RNA helicase domain of NAT10 shares many conserved characteristics with the DEAD box helicases^40,41^. These RNA helicases engage in several steps of RNA metabolism from splicing, nuclear export, translation, mRNA decay, to mRNA transport and storage. However, these RNA helicases interact with RNA in a sequence-independent manner^42^. Thus, in addition to further characterizing the detailed mechanisms utilized by helicase domain of NAT10, it will also be intriguing to characterize additional domains within NAT10 or protein binding partners of NAT10 that render its substrate specificity in future experiments. Given that NAT10 helicase activity regulates two fundamental catalyzing enzymes of glucose-derived serine/glycine biosynthesis pathway, limited serine and glycine in brain metastasis microenvironment may confer sensitivity to NAT10 helicase targeting agents.

## Methods

### Antibodies and chemicals

Antibodies used include anti-FLAG-M2 (Sigma, F1804), anti-α-tubulin (CST, 3873), anti-vinculin (CST, 13901; Sigma, v9131), anti-GAPDH (Abclonal, AC001), anti-NAT10 (Proteintech, 13365-1-AP; Bethyl, A304-385A), anti-ac4C (Abcam, ab252215), anti-cyclin D1 (CST, 2978), anti-V5 (Invitrogen, MA5-15253), anti-PHGDH (Proteintech, 14719-1-AP), anti-PSAT1 (Proteintech, 10501-1-AP), and anti-HSPA5 (Proteintech, 11587-1-AP). Chemicals used include Nocodazole (Sigma, M1404) and HA15 (MedChemExpress, HY-100437).

### ShRNA library and lentivirus for screen

Frozen bacterial stocks harboring the shRNA library were generated by the Thomas Westbrook lab at Baylor College of Medicine. pGIPZ plasmid harboring hairpins and barcodes were digested with Xho I and Mlu I and sub-cloned into the pINDUCER10 plasmid. The list of hairpin sequences is available in **Supplementary table 1**. For virus generation, HEK293T cells were transfected with 1.2 µg each of VSV-G, TAT, RAII, and HyPM packaging plasmids along with 11.2 µg of lentiviral plasmid. OptiMEM and TransIT-293 Transfection Reagent (Mirus, MIR2700) were used following manufacturer protocol. Viruses were collected at 48h and 72h, filtered through a 0.45 μm filter.

### Establishment of the *in vivo* selected MDA-MB-231-BrM3

The *in vivo* selected MDA-MB-231 brain metastasis derivatives cell line 231-BrM2 was previously obtained from Joan Massagué lab at the Sloan Kettering Institute at Memorial Sloan Kettering Cancer Center. To ensure its capability of colonizing the brain, 1×10^5^ 231-BrM2 cells were injected into nude mice intracardially. Brain metastases were monitored weekly with *in vivo* live imaging. At week five, the mice were euthanized, and the brain tissue was harvested and minced physically with razor blade and digested with medium containing 0.125% Type III Collagenase and 0.1% hyaluronidase for 1 hour at 37°C, and then with trypsin for another 15 min. After digestion, the tissue fragments were washed twice with 1×PBS and transferred to culture dish to sit down. Cells were split for a few passages until the fibroblasts were gone. The *in vivo* selected cell line, named MDA-MB-231-BrM3 (231-BrM3), was tested for GFP expression with FACS and tested for mycoplasma contamination. Finally, 231-BrM3 cells were re-injected into nude mice intracardially in parallel with 231-BrM2 cells to compare the brain colonization ability.

### Cell culture and media

MDA-MB-231, 231-BrM2, 231-BrM3, 231-BoM, 231-LM2, H2030-BrM3, MDA-MB-157, and HEK293T cells were cultured in Dulbecco’s Modified Eagle Medium (DMEM) supplemented with 10% (v/v) FBS (Gibco) and pen/strep. T47D was cultured in RPMI1640 supplemented with 10% FBS (Gibco) and pen/strep. Cells were periodically tested for mycoplasma contamination and authenticated using short tandem repeat profiling. For generation of cell lines for screening in 231-BrM3, viruses harboring pINDUCER10 constructs were titrated using the target cell lines, and cells were infected at an MOI of 1. 231-BrM3 stable cells were selected with 0.8 µg/mL puromycin.

DMEM-Like, CSF-like, w/o-Ser/Gly culture media were made by using DMEM without glucose, glutamine, serine, glycine, and sodium pyruvate (US Biological D9802-01), supplemented with 10% (v/v) inactivated dialyzed inactivated FBS (dIFS, Gibco, 26400044), pen/strep, GlutaMAX™ Supplement (2mM, Gibco, 35050061), D-glucose (4.5g/L); DMEM-Like medium contains 400μM serine and 400μM glycine, CSF-Like medium contains 25μM serine and 5μM glycine, and w/o-Ser/Gly does not contain serine and glycine. All pH-adjusted complete media were filtered through a 0.22-μm filter prior to use.

### Mini-pools generation for *in vitro* and *in vivo* screens

Mini-pools were created by equally mixing 11-13 individual inducible epigenetic regulator knockdown cell lines with shBUD31 and shCHEK1 as positive and negative controls, respectively. For *in vitro* screen, mini-pool cells were plated onto 10-cm dishes with or without 1 μg/mL of doxycycline. A portion of mini-pool cells were collected as day 0 samples as the controls. Every two days the cells were pelleted, and all samples were proceeded to gDNA isolation and gDNA qPCR. For *in vivo* screen, 5×10^5^ mini-pool cells were injected into nude mice through left ventricle for the brain screen. Brain metastases were monitored weekly with *in vivo* live imaging. At the end point, the mice were euthanized, and the targeted tissues were harvested for gDNA isolation and gDNA qPCR. For the screening readout analyses, all qPCR results were normalized to the value from day 0. The fold change was obtained from +/-doxycycline for both *in vitro* and *in vivo* screen.

### Animal studies

Six-week-old female, athymic mice (*Foxn1*^nu/nu^, NCI stock#553) were purchased from Charles River Laboratories for brain metastasis experiments with human cell lines. For *in vivo* screen, 5×10^5^ cells in 0.1 mL saline were injected intracardially. Mice under treated group were placed on doxycycline chow (Envigo, TD.01306) 5 days prior to injection. All the *in vivo* metastasis signals, including brain metastases and whole-body metastases, were monitored by weekly bioluminescence imaging with an IVIS system coupled to Living Image acquisition and analysis software (Xenogen). Luminescence signals were quantified at the indicated time points. Values of luminescence photon flux of each time point were normalized to the value obtained immediately after xenografting (day 0).

For brain colonization assay, cell lines were treated with doxycycline for 3 days prior to injection and 0.1 ml of PBS containing 20,000 shCtrl or shNAT10 231-BrM3 cells or 250,000 H2030-BrM3 were injected into the left ventricle of mice. Mice were placed on doxycycline chow (Envigo, TD.01306) 5 days prior to injections. Brain colonization was analyzed *in vivo* and *ex vivo* by BLI. Anesthetized mice (ketamine 100 mg/kg/ xylazine 10 mg/kg) were injected retro-orbitally with D-Luciferin (150 mg/kg) and imaged with an IVIS Spectrum Xenogen machine (Perkin Elmer). Bioluminescence analysis was performed using Living Image software.

For mammary fat pad tumor assays, Control and shNAT10 #1 231-BrM3 cells (1×10^6^) were resuspended in 0.1 ml of saline and Matrigel (Corning, 356231) mix, and then injected into mammary fat pad (the 4^th^ mammary glands) of NOD-SCID mice (6 weeks old). Tumor were monitored weekly by *in vivo* live imaging. Tumor volume at the end point was calculated by the measurements of tumor length (L) and width (W) as V=L×W^2^/2. Mice were euthanized when primary tumors reached 1,000 mm^3^. This work was performed according to NIH guidelines and all animal procedures were approved by the Institutional Animal Care and Use Committee of Yale University.

### Tissue harvest and gDNA isolation

Mice were euthanized and whole body perfused with 10 mL of PBS. For gDNA isolation, the harvested tissue was placed into a microcentrifuge tube and snap-frozen with liquid nitrogen. The frozen tissues were then placed into an aluminum block on dry ice. Each tube of the tissue was allowed to thaw enough for further mincing with surgical scissors, and then refrozen by dipping them in liquid nitrogen bath. This process was repeated 2-3 times until no visible tissue chunk was observed. Sixty mg of homogenized tissue was then aliquoted out and processed with the DNeasy Blood & Tissue Kits (Qiagen, 69504) following manufacturer’s protocols.

### Barcode gDNA qPCR

For barcode qPCR, isolated gDNA was diluted with water and Fast SYBR Green Master Mix (ThermoFisher, 4385614) and barcode primers were designed to amplify only one barcode sequence among the 100 unique barcodes in the entire library. The primer set targeting the TRE element in pINDUCER10 was used for normalization. The full list of barcode qPCR primers used for detection of hairpin abundance is available in **Supplementary table 2**.

### Histopathology

Mice were euthanized by CO_2_ asphyxiation and brain were harvested, immersion-fixed in 10 % neutral buffered formalin, processed, sectioned, and stained by hematoxylin and eosin (H&E) with routine methods by Yale Research Histology (Department of Pathology). Digital light microscopic images were recorded using a Keyence BZ-X700 immunofluorescent microscope.

### Knockdown and rescue cell lines

For cloning of the NAT10 WT and point mutants, primers flanking with EcoRI were designed against pICE-FLAG-NAT10-siR-WT and pICE-FLAG-NAT10-siR-G641E. Two-step PCR was performed to generate shRNA resistant mutants. The final PCR products were then cloned into pBABE-hygro plasmid by cut-and-paste method. The K290A and K426R mutants were further generated using site-directed mutagenesis.

For cloning of NAT10 WT and K290A into the inducible expression backbone, BP cloning primers were designed against pICE-FLAG-NAT10-siR-WT. The PCR product was then used for BP (Thermo Fisher, # 11789020) or LR (Thermo Fisher, 11791020) reaction into pDONR-211. The list of cloning oligos can be found in **Supplementary table 3**.

The recombinant lentivirus was produced by transient transfection in HEK293T cells with packing plasmid (psPAX2) and envelope VSV-G plasmid (pMD2.G) together with shRNA clones or overexpression vectors using Lipofectamine 2000 (ThermoFisher). Culture media were harvested 42-48 hours after transfection, filtered by syringe filter with 0.45 μm pore-size, and then added into the cells to be infected with 8 μg/mL polybrene. The infected cells were selected by specific antibiotics. Similarly, the recombinant retroviruses were produced by transient transfection of Phoenix Amphotropic (AMPHO) cells with pBABE-hygro expression vectors using Lipofectamine 2000 (ThermoFisher).

For *PHGDH*, *PSAT1*, and *HSPA5* knockdown 231-BrM3 cell lines, bacterial culture harboring pLKO.1 shRNA plasmids was purchased from Yale Cancer Center Functional Genomics Core or Sigma Predesigned shRNA. The extracted plasmid was transfected with packing plasmid (psPAX2) and envelope VSV-G plasmid (pMD2.G) using FuGENE® 6 Transfection Reagent (Promega, E2691). The following lentivirus harvest, infection, and selection steps were similar with the steps mentioned above. shRNA sequences are available in **Supplementary table 4**.

### Colony formation assay, WST-1 cell proliferation assay, and grow curve

Colony formation assays were done by seeding single cells in 6 or 12 well plates. Media was replenished every 3 days with indicated treatments. Colonies were fixed in 4% paraformaldehyde (PFA), followed by 0.5% crystal violet staining for 30 minutes at room temperature and rinsed with water. Quantification was performed using the ImageJ software plugin ColonyArea^43^. Statistical significance was determined using unpaired, two-tailed student’s t-test performed on intensity values from ColonyArea.

For WST-1 cell proliferation assay, cells were seeded at 1000 cells per well in 100 μl into 96 well white plates (Corning, 9154). 1-5 days after seeding as indicated, media was replicated by a 5% solution of WST-1 reagent (Roche, 11644807001) in media. The plate was incubated for 1 hour at 37 °C. The absorbance at 440 nm was measured, as well as the absorbance at 610 nm as a reference. The absorbance at 610 nm was subtracted from the 440 nm reading. The average of the media-only absorbance was subtracted from all wells. Unpaired, two-tailed student’s t-test was used to determine significance.

For growth curves, 10,000 cells per well in 1ml DMEM medium were seeded into 24 well plate (Costar, 3526) and 18 hours after seeding (defined as t = 0h), the media were replaced with the customized media and at least four pictures were taken for each well every 8 hours for 6 days with CellCyte X system. Confluency was calculated with the default analyzing software of CellCyte X system. For DOX inducible knockdown cell lines, DOX was added in DMEM medium to induce knockdown prior to the seeding and withdrawn in customized media during the proliferation assays.

### BrdU cell proliferation and cell cycle assays

Cell proliferation was assayed using Cell Proliferation ELISA BrdU kit (Roche, 11647229001). In brief, 3×10^3^ cells were seeded on 96 well plate and 5-Bromo-2′-deoxyuridine (BrdU) was added to culture medium at 18 hours after plating. BrdU labeling was proceeded for 1 hour before cell harvest. Absorbance at 450 nm was measured, as well as the absorbance at 690 nm as a reference. Unpaired, two-tailed student’s t-test was used to determine significance.

For cell cycle analysis, cells were treated with doxycycline for 3 days to induce shRNA expression then treated with nocodazole (40 ng/mL) in serum free medium for 24 hours for cell synchronization. Cells were then switched back to normal media. At 0, 9, 12, 15, 18, 24-hour time points, cells were collected and fixed in ice-cold 70% ethanol in 4°C overnight. Fixed cells were resuspended in PBS containing DAPI (1 μg/mL), and the analysis was performed on a FACS Stratedigm STD-13 cytometer.

### 3D organoid growth assay

Cells were resuspended in medium containing 2% FBS and 5% growth factor reduced Matrigel. Cells were then seeded into ultra-low attachment 24 well plate (Corning, 3473) at 1000 cells per well. Every 3 days, 50 µL of Matrigel containing medium were added to each well to refresh the cells. The spheroid growth was measured with Promega dual-luciferase reporter assay system (E1910). In brief, at day 12, spheroids were collected, pelleted, and lysed with 1× passive lysis buffer. Cell lysates were stored in - 80 °C at least overnight. At the day for luciferase assay, cell lysates were thawed and added with luciferase assay substrate, then the luciferase activity were measured immediately with plate reader.

### Cell migration and invasion assays

For Transwell migration and invasion assay, cell culture inserts (Corning, 353097) were placed in the 24 well plates. For Transwell migration assay, 2×10^4^ cells resuspended in medium containing 0.1% FBS were plated onto the upper chamber, and the medium containing 10% FBS was added to the lower chamber. Cells were incubated at 37°C for 5 hrs. For Transwell invasion assay, the upper side of the Transwell membrane was pre-coated with Matrigel (Corning, 354480). 5×10^4^ cells were plated onto the upper chamber as Migration assay. Cells were incubated at 37°C for 18 hrs. At the end point of incubated, cells that had migrated or invaded onto the lower membrane surface were fixed with 4% paraformaldehyde in PBS, stained with DAPI, and counted.

For wound healing assay, 5×10^5^ cells were seeded on 6 well plate and scratches were made using P200 tips; wash wells with 1×PBS and take pictures at t = 0h. Pictures were also taken at indicated time points and the area of the wound was calculated using ImageJ plugin *Wound_healing_size_tool*^44^.

### Apoptosis assays

Cells were treated with doxycycline for 3 days to induce shRNA expression prior to performing apoptosis assays. An additional plate of cells was treated with 50 μM of etoposide for 48 hours to induce apoptosis as a positive control. On the day of analysis, cells were harvested and stained with annexin V-Biotin (Biolegend, 640904) followed by Steptavidin-Cy7 (Biolegend, 405208) and helix NP Blue (Biolegend, 425305). Flow cytometry was performed to record the signals using allophycocyanin (APC) and Pacific Blue channels. Analyses were done using FlowJo software.

### Western blot

Cells were lysed in 1× high salt lysis buffer (50 mM Tris [pH=8], 320 mM NaCl, 0.1 mM EDTA, 0.5% NP-40, 10% glycerol) supplemented with 1× cOmplete protease inhibitor (Roche, 11836153001) and 1× PhosSTOP phosphatase inhibitor (Roche, 4906845001). Cell lysates were vortexed and centrifuged, the supernatants were subjected to protein quantification by Bradford reagent (Bio-Rad 5000006) and sample preparation by sample buffer (10% glycerol, 50 mM Tris-HCl [pH=6.8], 2% SDS, 0.01% bromophenol blue and 8% β-mercaptoethanol). Protein samples were resolved by SDS-PAGE according to standard protocol and transferred onto 0.45 μm nitrocellulose membranes (Bio-Rad, 1620115). The membranes were blocked with 1%∼5% milk in TBST (Tris buffered Saline with 0.1% Tween-20) at room temperature for at least 10 minutes. The membranes were incubated with primary antibody diluted in blocking condition overnight at 4°C. The nitrocellulose membranes were rinsed with 0.1% TBST three times for 5 minutes and incubated with indicated HRP-conjugated-secondary antibody at room temperature for 1 hour. Blots were rinsed with 0.1% TBST three times and visualized using Kwik-Quant system (Kindle Biosciences, D1001).

### RT-qPCR

Total RNA from cells was extracted using the RNeasy Plus Mini Kit (Qiagen, 74136) protocol. RNA was reverse-transcribed into cDNA using High-Capacity cDNA Reverse Transcription kit (ThermoFisher, 4385614). The resulting cDNA was diluted with DEPC water and Fast SYBR Green Master Mix (ThermoFisher, 4385614) was used for real-time PCR. GAPDH was utilized for normalization. Samples were run in quadruplicate and experiments were performed at least three times. Primer sequences are listed in **Supplementary table 5**. Unpaired, two-tailed student’s t-test was used to determine significance.

### RNA-sequencing

231-BrM3 cells from knockdown control or shNAT10 group were harvested with QIAzol Lysis Reagent (Qiagen) and homogenized using QIAshredder tubes (Qiagen). For each cell line, shRNA sequence was induced with doxycycline (1 μg/mL) for 3 days and 3 biological replicates were harvested at different passages. RNA isolation was performed using miRNeasy with on-column DNase digestion. ERCC spike-in RNA was added in proportion to the number of cells obtained during cell counts. Library generation was performed using TruSeq stranded mRNA library prep kit (Illumina). Paired-end sequencing was performed using an Illumina HiSeq4000 sequencer, generating an average of 59 million reads per library. Reads were aligned to hg38 and gene counts to GENCODEv96 transcripts were obtained using STAR aligner v2.7.0 with default parameters. The hg38 and GENCODEv96 annotations were appended to include the ERCC sequences. DESeq2 was used to obtain differential gene expression, and HTSFilter was used to filter for expressed genes. Significant differences were identified using a Benjamini-Hochberg adjusted p-value cut-off of 0.05. Differentially expressed gene list is available in **Supplementary table 6**. The GO analysis of differential expressed genes was performed with The Database for Annotation, Visualization and Integrated Discovery (DAVID, https://david.ncifcrf.gov/)^45^ based on Knowledgebase v2024q1.

### DIA-mass spectrometry

shCtrl and NAT10 knockdown (shNAT10 #1) 231-BrM3 cells were induced with doxycycline (1 μg/mL) for 3 days and 3 biological replicates were harvested at different passages. The protein extraction, digestion, DIA-MS, and data analysis were described in Mehnert et. al^29^. Protein list of DIA-MS is available in **Supplementary table 7**. The GO analysis of differential expressed proteins was performed with DAVID (https://david.ncifcrf.gov/) based on Knowledgebase v2024q1.

### Immuno-northern blot

Equal amounts of total RNA (15 μg) or poly(A) RNA were mixed with 4 times volume of RNA sample buffer (65% formamide, 22% formalin, 13% 10× MOPS, ethidium bromide 8 μg/mL) and 1 time volume of loading dye (50% glycerol, 1 mM EDTA, 0.3% each bromophenol blue, 0.3% xylene cyanol), heated to 65°C for 15 min and separated on 1% agarose denaturing gel. Loading control was verified by UV imaging before transfer. RNA was transferred onto Amersham Hybond-N+ membranes (GE Healthcare) by capillary transfer using 1X transfer buffer (3M NaCl, 0.01N NaOH) for 1.5 hr. Membranes were rinsed with 6× SSC buffer, crosslinked twice with 150 mJ/cm^2^ in the UV254nm Stratalinker 2400, blocked with 5% non-fat milk in 0.1% Tween-20 PBS (PBST) for 30 minutes at room temperature, and probed overnight with anti-ac4C antibody in 1% non-fat milk (1 : 10000) at 4°C. Membranes were next washed three times with 0.1% PBST, incubated with HRP-conjugated secondary anti-rabbit IgG in 1% non-fat milk at 4°C overnight, washed three times with 0.1% PBST and developed with the SuperSignal ELISA Femto Maximum Sensitivity Substrate (ThermoFisher Scientific). Chemiluminescence was detected using Kwik-Quant Imager Camera.

### Translation rate assay

Cells were starved of L-methionine for 30 minutes and subsequently incubated with 50 μM homopropargylglycine (HPG; Life Technologies, C10186) for 1 to 4 hours in treatment media. Cells were then trypsinized and fixed in 4% paraformaldehyde. A Click-IT kit (Life Technologies, C10269) was used to label HPG. Labeled cells were analyzed using a CytoFLEX flow cytometer. Translation rates were determined based on the slope of HPG incorporation over time.

### Statistical analysis

Comparisons between two groups were performed using an unpaired two-side student’s t test unless indicated otherwise. Graphs represent either group mean values ± SEM or individual values (as indicated in the figure legends). For animal experiments, each tumor graft was an independent sample. All experiments were reproduced at least three times.

## Data availability

All data used to generate the results are found in the extended data and supplementary information. RNA-seq data generated by the current study have been deposited into the National Center for Biotechnology Information (NCBI) Gene Expression Omnibus database under GSE269160 (reviewer token: uribciuslliftyx). Other publicly available datasets used in the study include GSE138122, GSE148005, GSE183862, and The AURORA US Metastasis Project (GSE193103).

## Code availability

The codes for the analysis of NAT10 and its targets expression in MDA-MB-231 and 231-BrM3 have been deposited in GitHub (https://github.com/wesleylcai/bmcmedgenomics2020_metastasis).

## Acknowledgements

We would like to thank all members from Drs. Yan and Nguyen’s laboratory at Yale School of Medicine for helpful discussions, Dr. Mei Zhong at Yale Stem Cell Center Genomics Core facility for helping with sample preparation for RNA-seq, Dr. Joan Massagué at Memorial Sloan Kettering Cancer Center for providing MDA-MB-231, 231-LM2, 231-BrM2, and 231-BoM cells, and Dr. Narendra Wajapeyee at the University of Alabama Birmingham for helping with compiling the epigenetic gene list. RNA sequencing at Yale Stem Cell Center Genomics Core facility was supported by the Connecticut Regenerative Medicine Research Fund and the Li Ka Shing Foundation. This work is supported by the Department of Defense Breast Cancer Research Program Awards W81XWH-15-1-0117 and W81XWH-21-1-0411 (to Q.Y.), National Institutes of Health Awards R01CA237586 (to QY), R01CA166376 (to DXN), F31CA243295 (to J.F.C.), and P30CA016359 (to the Yale Comprehensive Cancer Center), Yale Cancer Center Class of ’61 Cancer Research Award (to QY and DXN), Leslie H. Warner Postdoctoral Fellowship (to P.X.), NSF Graduate Research Fellowship DGE-1122492 (to W.L.C.), and CSC fellowship (to H.C.). The funders played no role in the design of the study and collection, analysis, and interpretation of data and in writing the manuscript.

## Author contributions

Q.Y. and D.X.N. conceived the study; Q.Y. oversaw the project. J.F.C. and W.L.C. designed and carried out the screens with the help from T.F.W.; J.F.C. designed and performed most in vitro and in vivo experiments with help from W.L.C., H.C., G.B., M.Z., A.B., E.D.K., Y.Z., M.Y., T.T., J.L.M., and S.H.; P.X. finished the identification of downstream effectors of NAT10 and performed mechanistic investigation; E.W. and D.X.N. performed the lung cancer brain metastasis experiments and analysis; W.L.C. performed the RNA-Seq analysis; W.L. and Y.L. performed DIA-MS experiment and analysis; P.X. performed the analysis of public datasets and integration of datasets; P.X., J.F.C. and Q.Y. wrote and revised the manuscript, and J.L.M. and S.H. revised the manuscript.

## Competing interests

Q. Y. and D.X.N. have received research funding unrelated to this study from Astra Zeneca Inc. Q.Y. is a member of Scientific Advisory Board of AccuraGen Inc. The other authors declare no competing interests.

---

**Extended Data Fig. 1.**
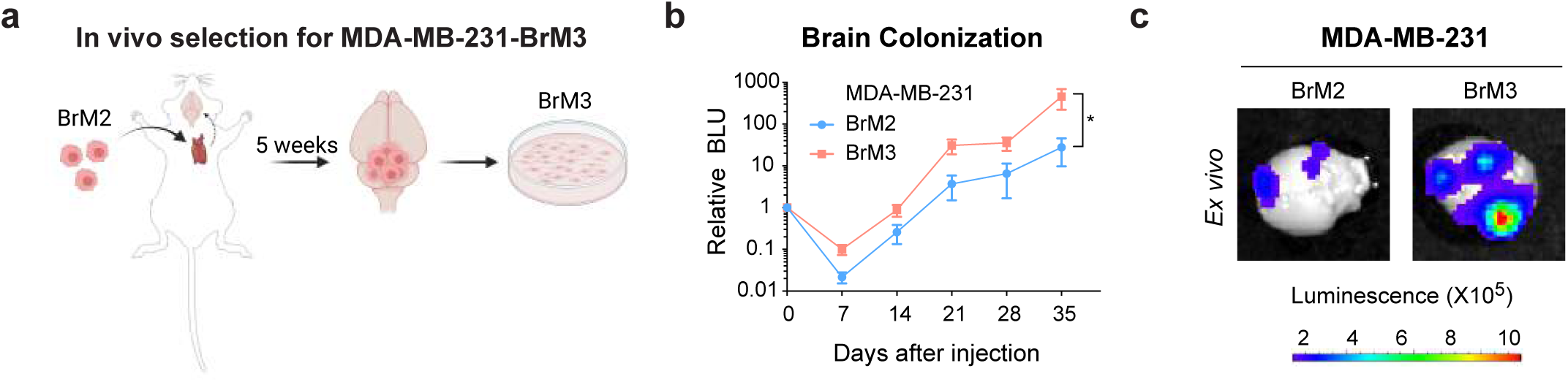
| Generation of triple-negative breast cancer cell line with brain organotropism. **a**, Schematic of *in vivo* selection process to generate MDA-MB-231-BrM3 (231-BrM3) cell line. **b**, Normalized bioluminescence signals of brain metastases of mice injected intracardiacally with MDA-MB-231-BrM2 (231-BrM2) or 231-BrM3 cells from (**a**). The data represent average ± SEM. Significance was determined using two-tailed student’s t-test. *, *P* < 0.05. **c**, Representative bioluminescence images of mice brain metastases in (**a**) at day 35.

**Extended Data Fig. 2.**
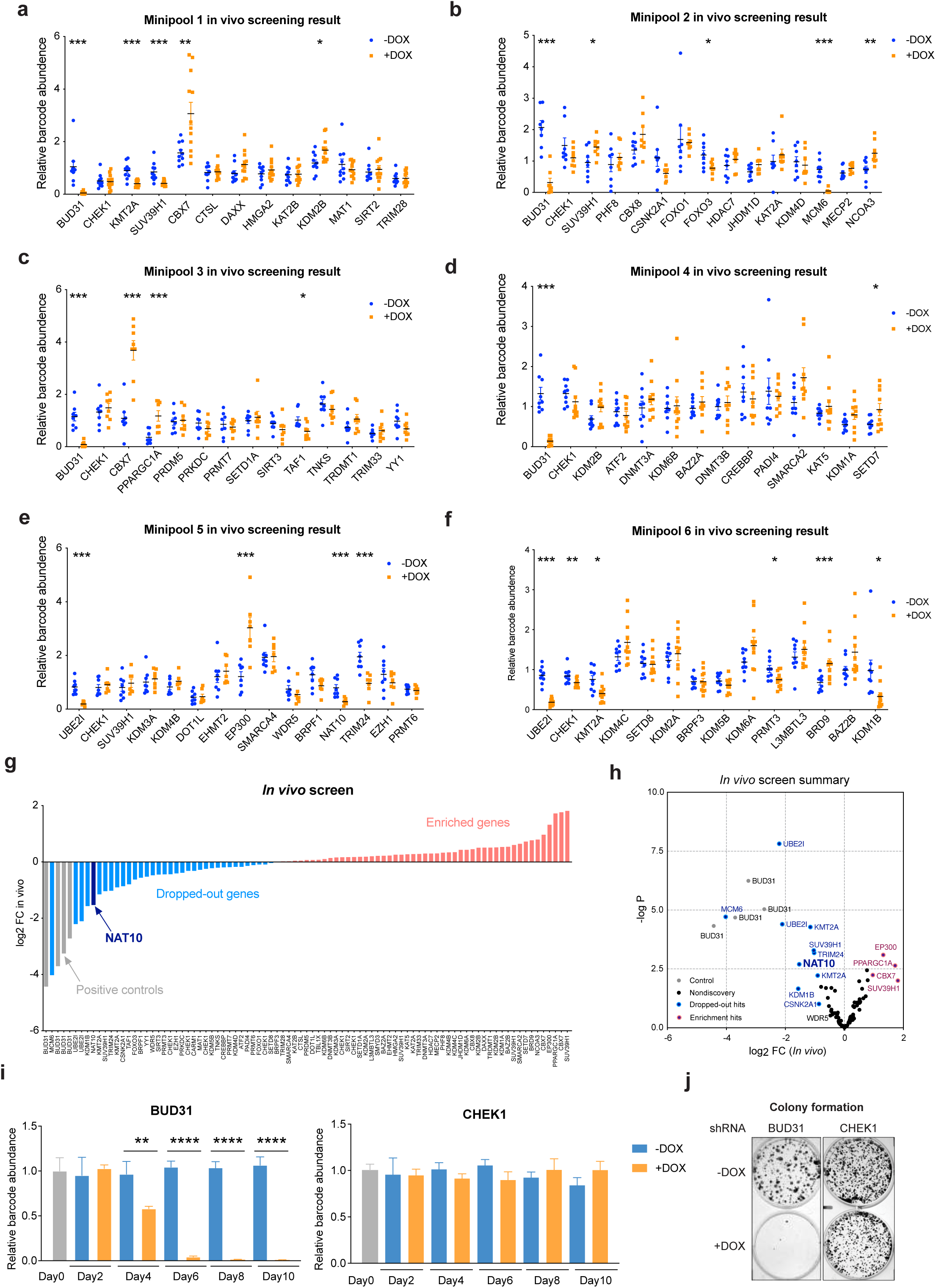
| Screening results of breast cancer brain metastasis. **a-f**, Relative abundance of barcode for shRNA against each epigenetic factor in the library in brain tissues from control and doxycycline-treated mice. **g**, Waterfall plot of the *in vivo* screening results. Log_2_ (fold change) of each cell line was listed in ascending. **h**, Volcano plot demonstrates the results of the *in vivo* screen. Each data point is an average of 10-20 mice. Discovery hits: *P* < 0.05 and log_2_FC > 0.8 or log_2_FC < -0.8. **i**, Relative abundance of 231-BrM3 cells stably expressing shRNA against *BUD31* and *CHEK1* after the indicated days of *in vitro* culture under control or doxycycline (1 µg/mL) treatment. Data normalized to abundance at the time of mini-pool mixture (Day 0). **j**, Colony formation assays of 231-BrM3 cells after 9 days of either control or doxycycline (1 µg/mL) treatment. Significance was determined in (**a-f**) and (**i**) using unpaired student’s t-test. *, *P* < 0.05; **, *P* < 0.01; ***, *P* < 0.001; ****, *P* < 0.0001.

**Extended Data Fig. 3.**
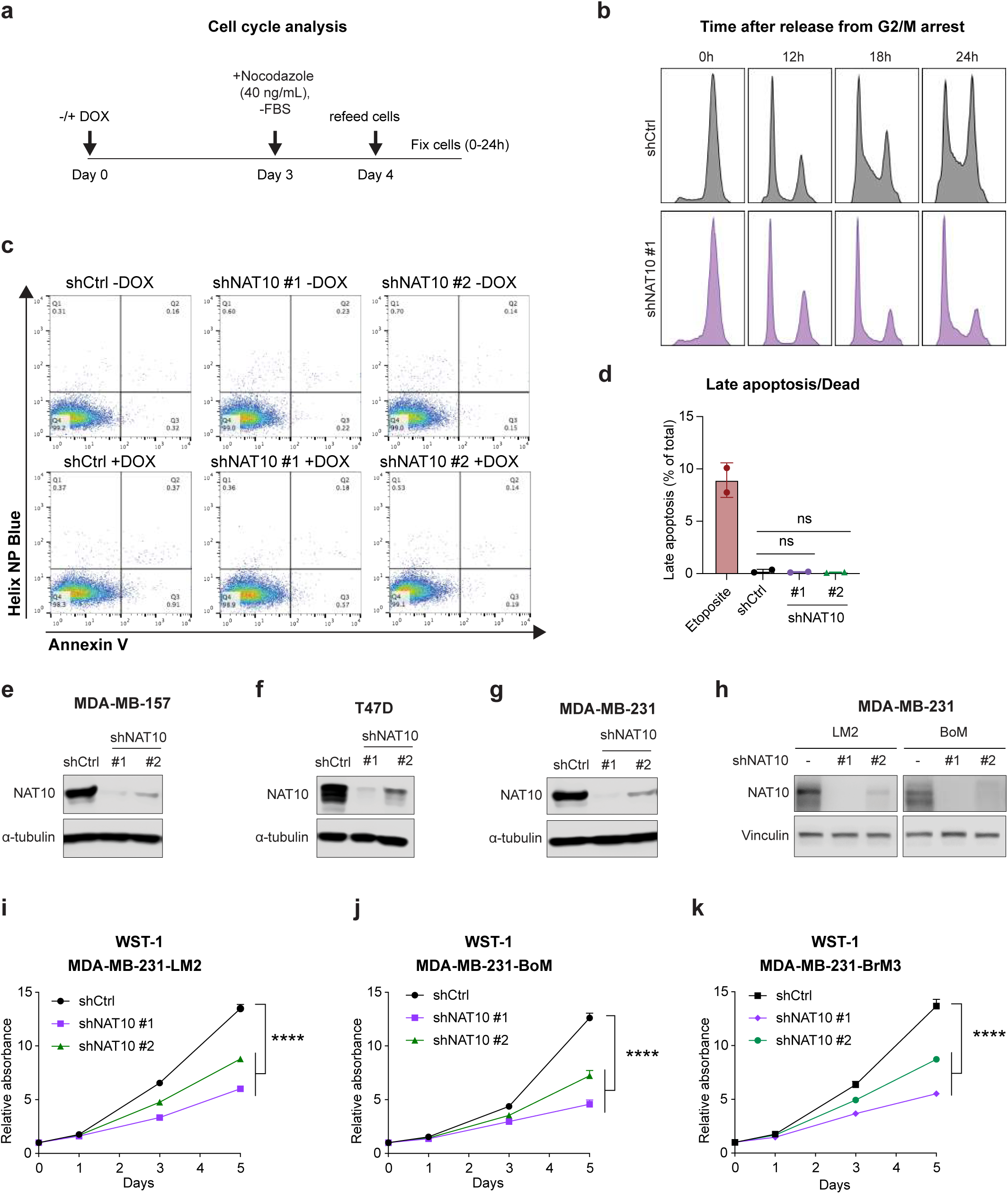
| NAT10 silencing disrupts cell cycle and attenuates proliferation of breast cancer cells with distinct metastatic organotropisms. **a**, Schematic of cell cycle analysis using the synchronizing method of Nocodazole (40 ng/mL) in combination with serum starvation for 24 hours. **b**, Time after release from G2/M arrest in 231-BrM3 cells. **c**, Apoptosis assay determined by Helix NP-Blue and Annexin V staining. 231-BrM3 cells from (b) were cultured with regular media with control or doxycycline (1 µg/mL) induction for 3 days. Representative experiment was shown; **d**, Late apoptosis versus dead cell ratio treated with etoposide and doxycycline; e-h, Western blots of NAT10, α-tubulin or Vinculin in indicated cell lines harboring inducible control or NAT10 targeting shRNAs (shNAT10 #1 and shNAT10 #2) after 3 days of doxycycline (1 µg/mL) induction. **i-k**, WST-1 proliferation assays of MDA-MB-231 organotropic derivatives after indicated days of doxycycline (1 µg/mL) treatment. Significance in (d and i-lk) was determined using unpaired student’s t-test. ns, not significant; ****, *P* < 0.0001.

**Extended Data Fig4.**
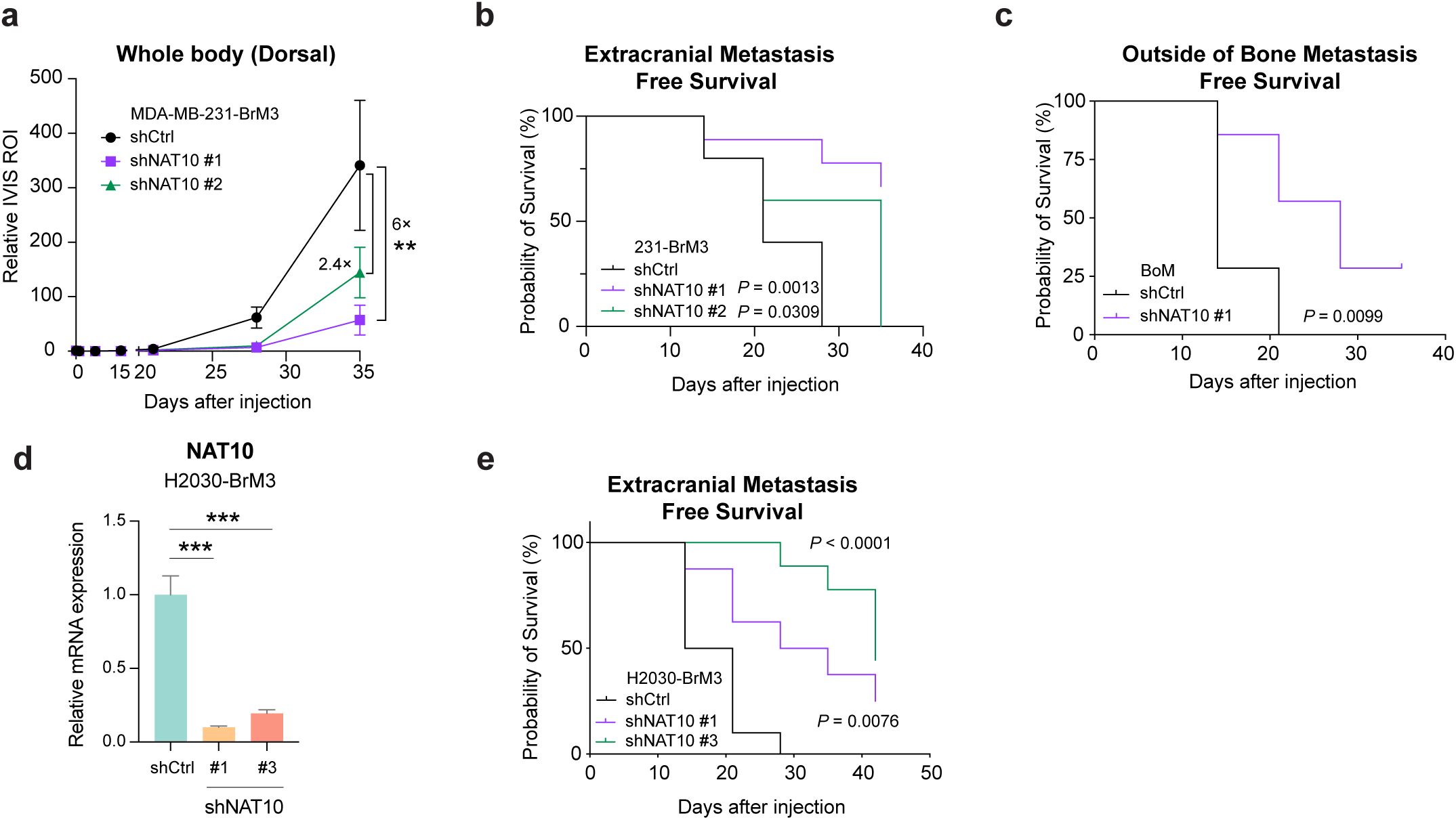
| NAT10 is an essential factor in multiple metastasis settings. **a**, Normalized bioluminescence signals of whole-body metastasis of mice injected intracardially with 231-BrM3 cells harboring inducible control, shNAT10 #1, or shNAT10 #2 and kept under doxycycline chow. The data represent average ± SEM (Unpaired student’s t-test; **, *P* < 0.01). **b**, Kaplan-Meier plot of extracranial metastasis-free survival for mice described in Fig. 3e. shCtrl (n=10), shNAT10 #1 (n=9), and shNAT10 #2 (n=10) were analyzed (Log rank Mantel-Cox test). **c**, Kaplan-Meier plot of metastasis-free survival outside of bone for mice described in Fig. 3k (Log rank Mantel-Cox test). **d,** RT-qPCR analysis of *NAT10* showing the knockdown effects of shRNA targeting NAT10 in H2030-BrM3 cells. **e**, Kaplan-Meier plot of whole-body metastasis-free survival in Fig. 3o. (Log rank Mantel-Cox test).

**Extended Data Fig. 5.**
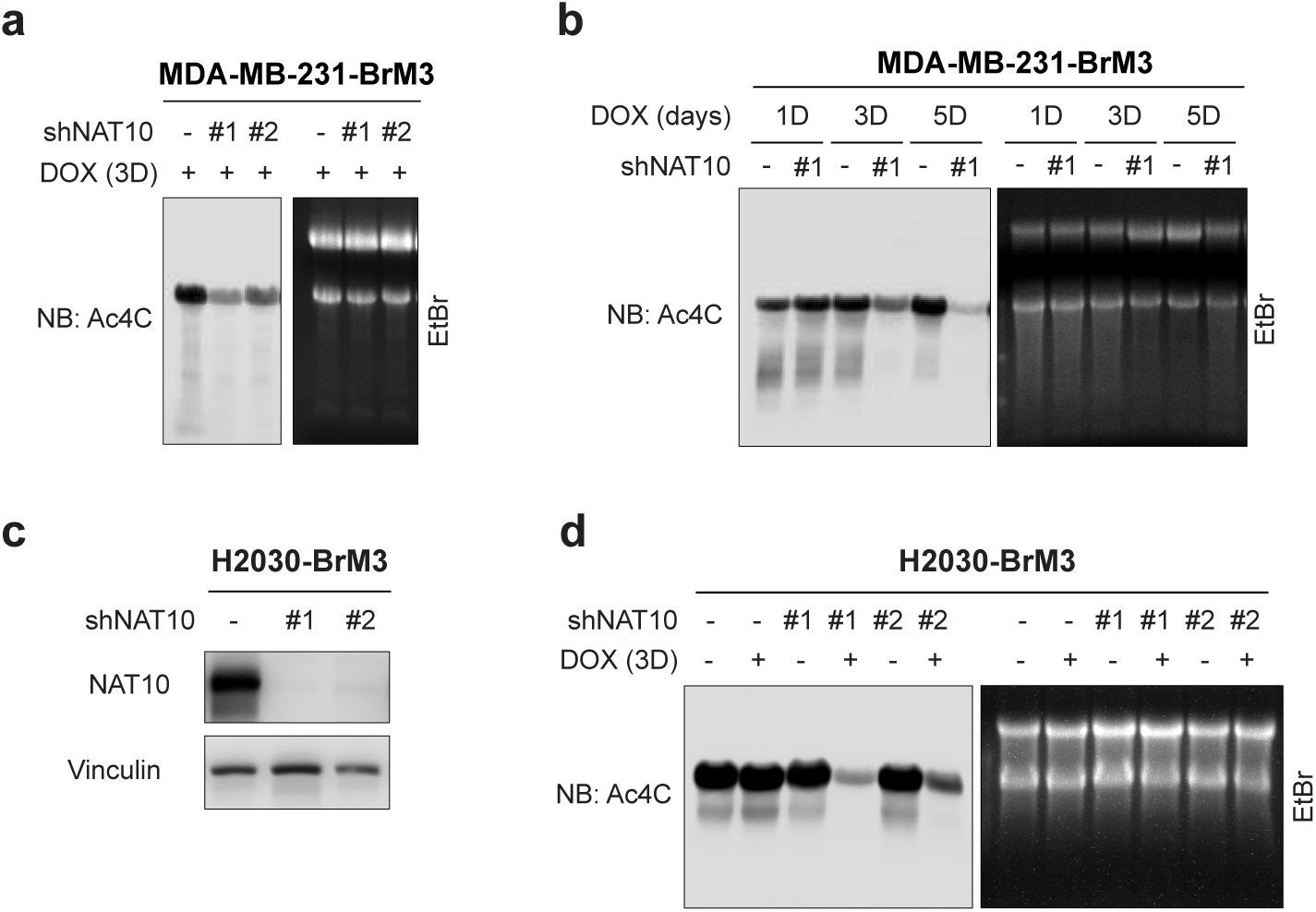
| N-acetyltransferase function of NAT10 is active in brain metastatic derivatives. **a**, Immuno-Northern blots of ac4C modification in total RNA extracted from 231-BrM3 cells harboring inducible control or NAT10-targeting shRNAs after 3 days of doxycycline (1 μg/mL) induction. **b**, Immuno-Northern blots of ac4C modification in total RNA extracted from 231-BrM3 cells harboring inducible control or NAT10-targeting shRNA after doxycycline (1 μg/mL) induction for 1, 3, or 5 consecutive days. **c**, Western blot analysis of NAT10 and vinculin showing the knockdown effects of shRNA targeting NAT10 in H2030-BrM3 cells. **d**, Immuno-Northern blots of ac4C modification in total RNA extracted from H2030-BrM3 cells harboring inducible control or NAT10-targeting shRNAs after 3 days of control or doxycycline (1 μg/mL) induction. In (**a**, **b, d**), denatured agarose gels were stained with EtBr as loading control on the right, in which the upper and lower band were 28S and 18S rRNA, respectively.

**Extended Data Fig. 6.**
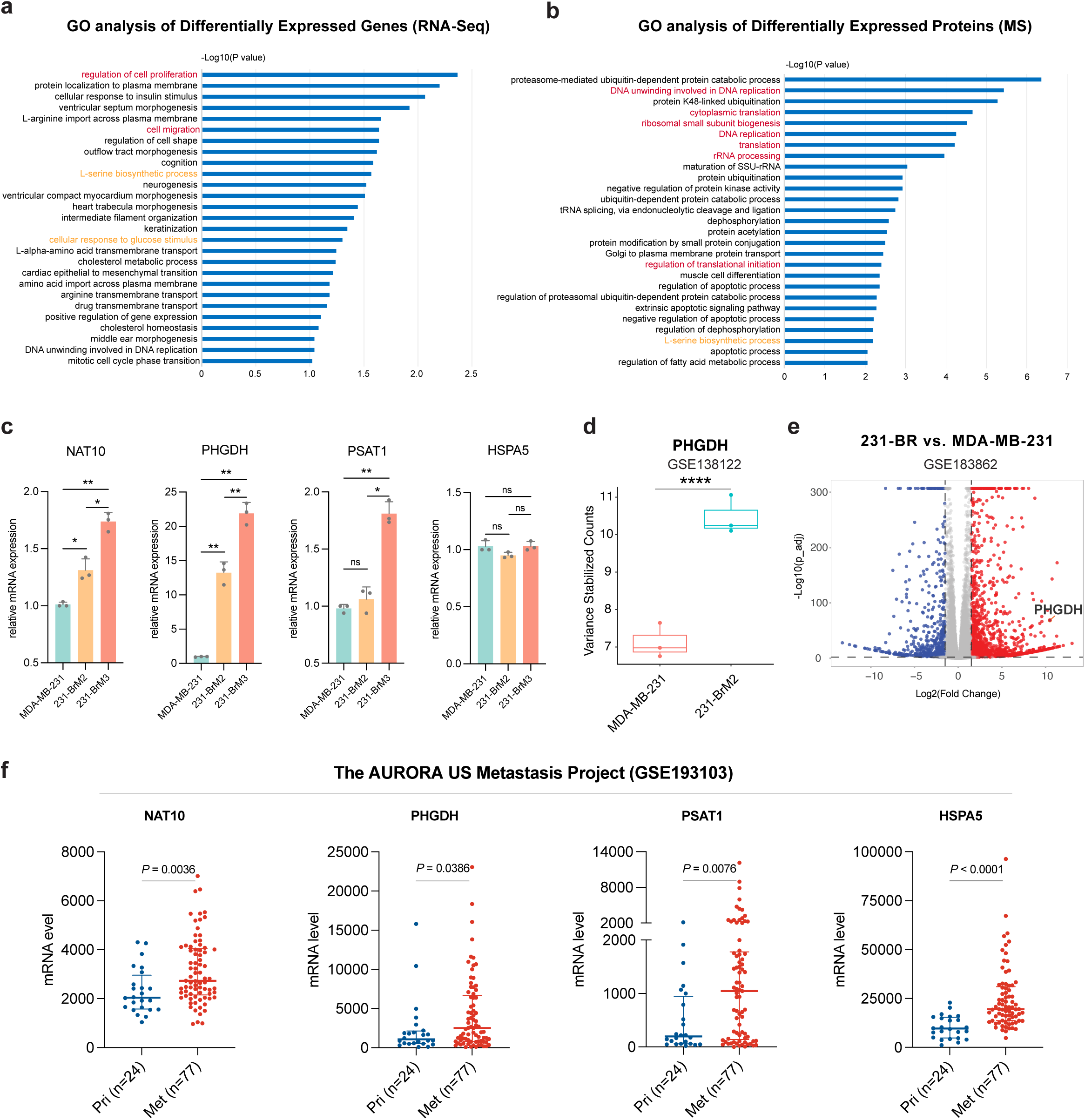
| NAT10 and its targets PHGDH, PSAT1, HSPA5 are highly expressed in metastatic breast cancer cells. **a-b**, The enriched Biological Process (BP) terms of differentially expressed genes/proteins in RNA-Seq (**a**)/DIA-MS (**b**) of NAT10 knockdown (shNAT10 #1) versus control (shCtrl) 231-BrM3 cells. For both RNA-Seq and DIA-MS, the differentially expressed candidates were defined as *P* < 0.5 and log_2_ (Fold Change) > 0.3 or < -0.3. Terms marked in red indicate the conventional biological processed that NAT10 involves, while terms marked in orange indicate the glucose-derived serine/glycine biosynthesis pathway. **c**, Relative mRNA levels of NAT10, PHGDH, PSAT1, and HSPA5 in parental MDA-MB-231, 231-BrM2, and 231-BrM3 cell lines. **d**, mRNA level (normalized counts) of PHGDH in parental MDA-MB-231 and 231-BrM2 from our previously published dataset (GSE138122). **e**, Volcano plot of the differentially expressed genes in MDA-MB-231 brain metastasis variant (231-BR) versus MDA-MB-231 from a publicly available RNA-seq dataset (GSE183862). **f**, NAT10, PHGDH, PSAT1, HSPA5 mRNA levels in primary breast tumors and their matched metastases in The AURORA US Metastasis Project (GSE193103). Met, metastases. Unpaired student’s t-test was used in (**c**, **d**), while paired student’s t-test was used in (**f**). *, *P* < 0.05; **, *P* < 0.01; ***, *P* < 0.001.

**Extended Data Fig. 7.**
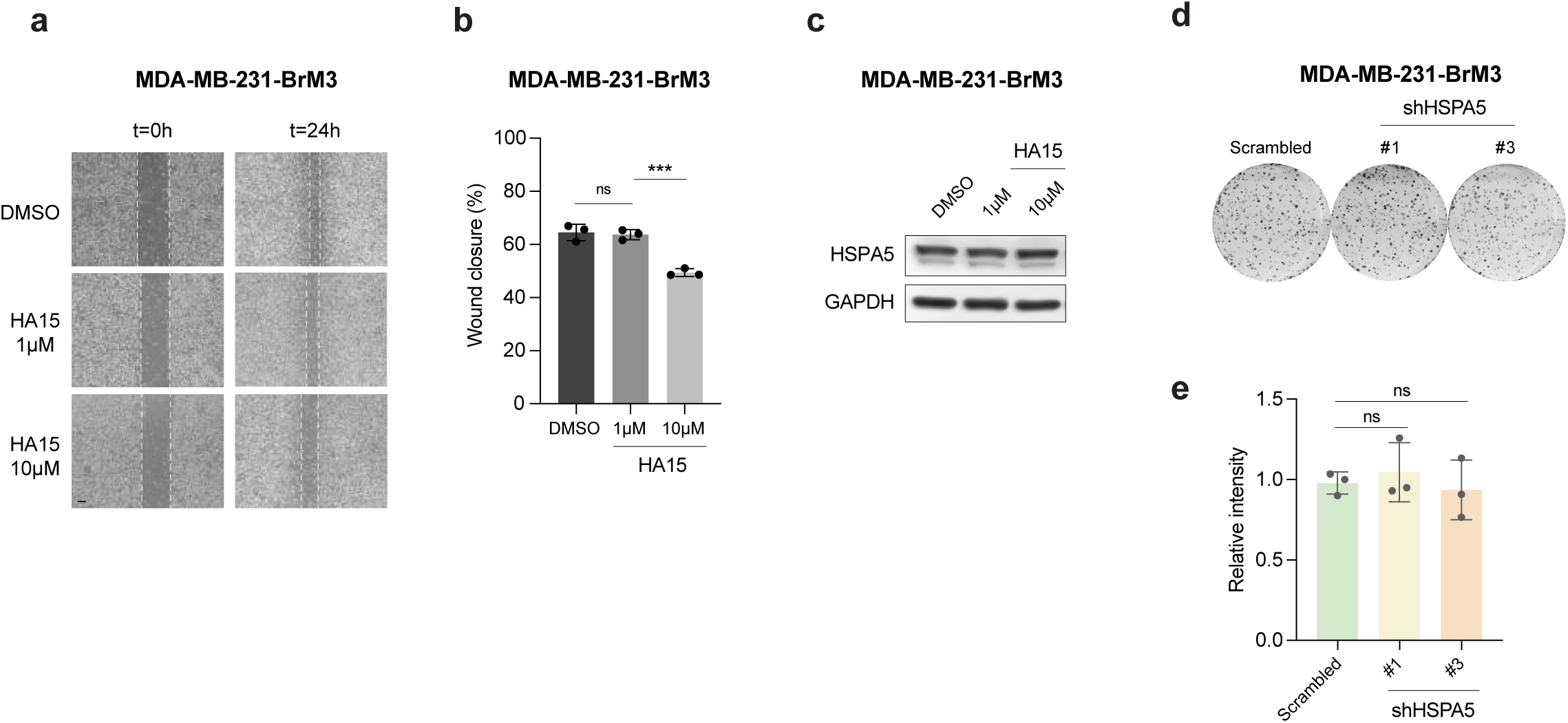
| NAT10-regulated HSPA5 modulates the migration of metastatic breast cancer cells. **a-b**, Wound healing assay of 231-BrM3 treated with HA15, a HSPA5 inhibitor. Representative pictures (**a**) and quantification (**b**) were shown. **c**, The protein level of HSPA5 in 231-BrM3 treated with increasing concentration of HA15. **d-e**, Colony formation assay of 231-BrM3 cells with or without HSPA5 knockdown. Representative pictures (**d**) and quantification (**e**) were shown. Unpaired student’s t-test was used in (**b** and **e**). ns, not significant; ***, *P* < 0.001.

## Supplementary Table list

Supplementary Table 1: The list of hairpin sequences for screens.

Supplementary Table 2: The full list of barcode qPCR primers for hairpin abundance detection.

Supplementary Table 3: The list of cloning oligos.

Supplementary Table 4: The sequences of shRNAs used in functional characterization.

Supplementary Table 5: The list of qPCR primers.

Supplementary Table 6: Differentially expressed genes in 231-BrM3 cells with shNAT10 #1 versus 231-BrM3 cells with shCtrl.

Supplementary Table 7: Differentially expressed proteins in 231-BrM3 cells with shNAT10 #1 versus 231-BrM3 cells with shCtrl.

## Notes

### Competing Interest Statement

Q. Y. and D.X.N. have received research funding unrelated to this study from Astra Zeneca Inc. Q.Y. is a member of the Scientific Advisory Board of AccuraGen Inc. The other authors declare no competing interests.

